# Covalent macrocyclic proteasome inhibitors mitigate resistance in *Plasmodium falciparum*

**DOI:** 10.1101/2023.07.03.547579

**Authors:** John M. Bennett, Kurt E. Ward, Ryan Muir, Stephanie Kabeche, Euna Yoo, Tomas Yeo, Grace Lam, Hao Zhang, Jehad Almaliti, Gabriel Berger, Franco F. Faucher, Gang Lin, William H. Gerwick, Ellen Yeh, David A. Fidock, Matthew Bogyo

**Affiliations:** Department of Chemistry, Stanford University, Stanford, CA, USA; Department of Microbiology and Immunology, Columbia University Medical Center, New York, NY, USA; Center for Malaria Therapeutics and Antimicrobial Resistance, Columbia University Medical Center, New York, NY, USA; Department of Pathology, Stanford University School of Medicine, Stanford, CA, USA; Department of Biochemistry, Stanford University School of Medicine, Stanford, CA, USA; Chemical Biology Laboratory, Center for Cancer Research, National Cancer Institute, National Institutes of Health, Frederick, MD, USA; Department of Medicine, Division of Gastroenterology and Hepatology, Stanford University School of Medicine, Stanford, CA, USA; Department of Microbiology and Immunology, Weill Cornell Medicine, New York, NY, USA; Scripps Institution of Oceanography, University of California San Diego, La Jolla, CA, USA; Skaggs School of Pharmacy and Pharmaceutical Sciences, University of California San Diego, La Jolla, CA, USA; Department of Microbiology and Immunology, Stanford University School of Medicine, Stanford, CA, USA; Division of Infectious Diseases, Columbia University Medical Center, New York, NY 10032 USA

## Abstract

The *Plasmodium* proteasome is a promising antimalarial drug target due to its essential role in all parasite lifecycle stages. Furthermore, proteasome inhibitors have synergistic effects when combined with current first-line artemisinins. Linear peptides that covalently inhibit the proteasome are effective at killing parasites and have a low propensity for inducing resistance. However, these scaffolds generally suffer from poor pharmacokinetics and bioavailability. Here we describe the development of covalent, irreversible macrocyclic inhibitors of the *P. falciparum* proteasome. We identified compounds with excellent potency and low cytotoxicity, however, the first generation suffered from poor microsomal stability. Further optimization of an existing macrocyclic scaffold resulted in an irreversible covalent inhibitor carrying a vinyl sulfone electrophile that retained high potency, low cytotoxicity, and had acceptable metabolic stability. Importantly, unlike the parent reversible inhibitor that selected for multiple mutations in the proteasome, with one resulting in a 5,000-fold loss of potency, the irreversible analog only showed a 5-fold loss in potency for any single point mutation. Furthermore, an epoxyketone analog of the same scaffold retained potency against a panel of known proteasome mutants. These results confirm that macrocycles are optimal scaffolds to target the malarial proteasome and that the use of a covalent electrophile can greatly reduce the ability of the parasite to generate drug resistance mutations.

## INTRODUCTION

*Plasmodium falciparum* causes the most severe and lethal cases of malaria in humans. The disease remains a threat to 40% of the world’s population and causes over 600,000 deaths per year, mainly among children under the age of five.^1^ The emergence of resistance to the current frontline drug artemisinin (ART) and its partner drugs in ART-based combination therapies (ACTs) is particularly concerning, because these drugs are widely used in endemic populations.^2–6^ There is an urgent need to develop drugs with novel mechanisms of action and broad therapeutic potential to improve interventions and overcome multidrug resistance.

An attractive target for new drugs is the *Plasmodium* proteasome, which is essential throughout the parasite life cycle. Inhibitors of the proteasome synergize with ART to enhance killing of the parasite *in vivo*.^7,8^ A significant challenge for the development of *Plasmodium* proteasome inhibitors is optimizing selectivity over the human enzyme. We have previously used substrate screening and cryo-electron microscopy to evaluate differences in ligand binding preferences between the human and parasite proteasomes.^9^ This has enabled the discovery of potent, irreversible covalent inhibitors that are selective for the *Plasmodium* proteasome (**Figure 1**). These selective inhibitors enabled validation of the proteasome as a viable anti-malarial target using mouse models of infection.^7^ Building on this foundation, we and others developed various peptidic scaffolds and incorporated non-natural amino acids to increase overall selectivity and potency for the malaria proteasome.^10–12^ These parasite-selective compounds have enabled more detailed studies of the parasite’s potential to generate resistance mechanisms against proteasome inhibitors.^13^ Though promising, linear peptide inhibitors suffer from poor bioavailability and stability *in vivo*.^11^

**Figure 1.**
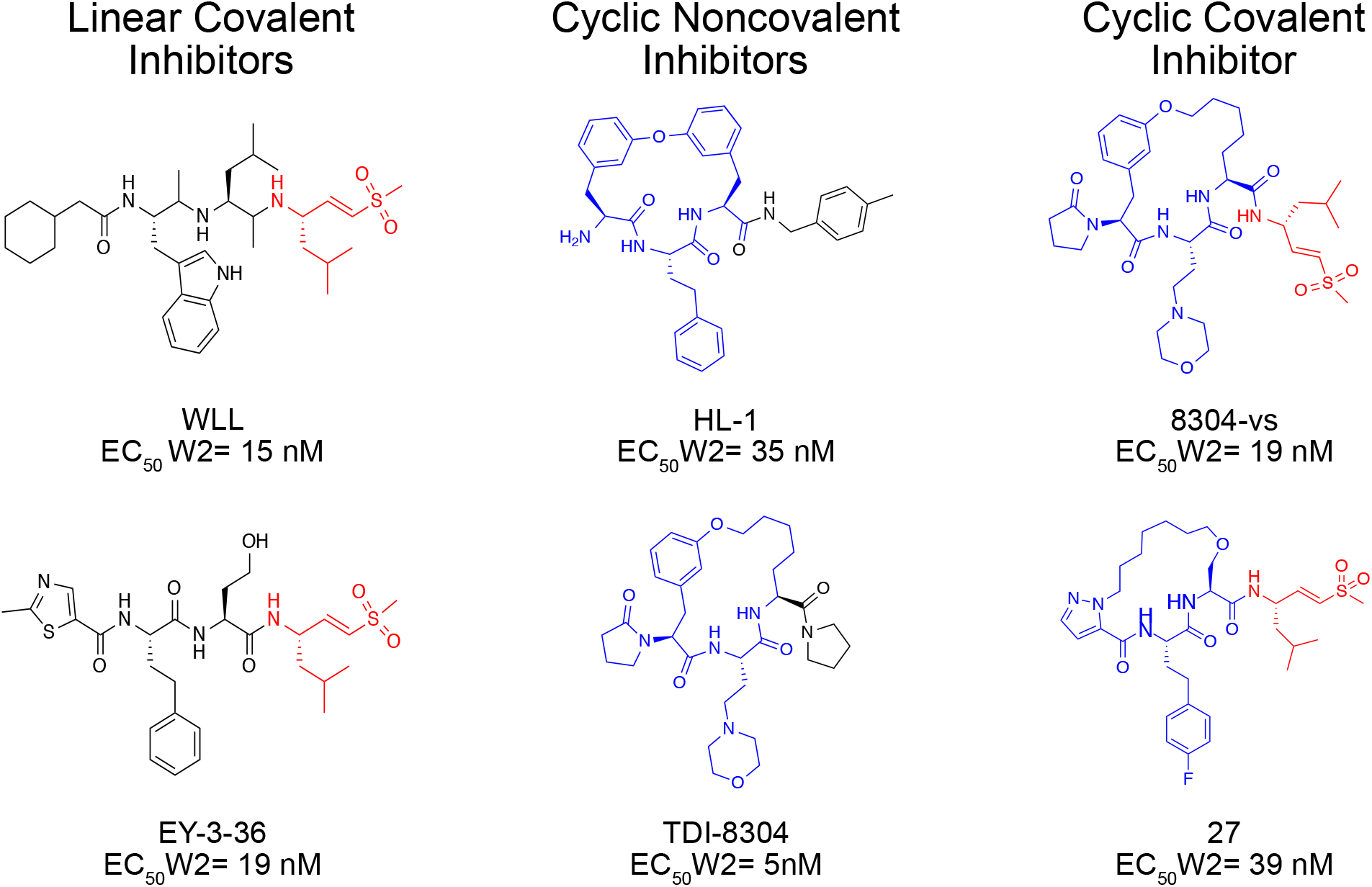
Chemical structures of representative macrocyclic covalent proteasome inhibitors are shown on the right. Previously reported linear covalent or noncovalent macrocyclic inhibitors are shown on the left as a reference.^9,11,16,17^ The covalent electrophilic warhead leucine vinyl sulfone is colored in red on each structure and the macrocycles are colored blue. EC_50_ values represent the mean concentration required to inhibit *P. falciparum* asexual blood stave parasite growth by 50%.

Peptide macrocycles have the potential to overcome many of the limitations of linear peptides as therapeutic agents.^14^ The former are generally more stable than their linear counterparts because cyclization drastically reduces the number of rotatable bonds, can mask hydrogen bond donors through internal hydrogen bonding, and can reduce proteolytic metabolism.^15^ Our prior screen of a small library of reversibly binding proteasome inhibitors identified a macrocyclic peptide containing a biphenyl ether with low nanomolar potency against intra-erythrocytic parasites and low toxicity in human cells.^16^ However, this molecule had poor solubility and rapid clearance. This scaffold was further developed to increase its potency and to optimize its pharmacokinetic properties, resulting in the promising lead molecule TDI-8304.^17,18^ However, we recently observed that this class of noncovalent cyclic peptide inhibitors can induce relatively high levels of resistance compared to similar classes of linear covalent irreversible proteasome inhibitors.^19^

We set out to assess the potential of various macrocyclic scaffolds to be potent, selective, and metabolically stable proteasome inhibitors. We also sought to evaluate how switching from reversible to covalent impacts resistance generation in the parasite. We describe herein the synthesis and characterization of a series of covalent macrocyclic inhibitors that explore the effects of ring size, capping elements, use of non-natural amino acids and overall hydrophobicity on potency, selectivity, and metabolic stability. While many of the compounds are potent and selective inhibitors of *Plasmodium* growth, some of the most potent molecules suffered from instability in both mouse and human microsomes. Therefore, we synthesized a covalent irreversible analog of the optimized macrocyclic scaffold TDI-8304 by adding a vinyl sulfone electrophile (8304-vs). Both TDI-8304 and 8304-vs showed selective and potent inhibition of the parasite and favorable metabolic stability with exceptionally low toxicity to host cells. Resistance studies demonstrated that both compounds induced a similar point mutation in the proteasome that altered the inhibitory potency of the molecules. However, the covalent binding version of the compound only encountered mild resistance with a 5-fold drop in potency, whereas the reversible binding molecule had a 5,000 drop in potency for this same proteasome mutant. We further assessed the importance of covalency by synthesizing an epoxyketone analog of TDI-8304 (8304-epoxy) and demonstrated that both the vinyl sulfone and the epoxyketone derivatives showed low cross resistance with known proteasome mutations. These results suggest that cyclic peptides are potentially optimal scaffolds for proteasome inhibitors and that use of covalent binding functional groups can help to reduce resistance liabilities.

## RESULTS

### Structure-Activity Relationship

Previously we used substrate profiling methods to define the specificity of the parasite proteasome relative to the human counterpart.^11^ We determined that a bulky aromatic group at the P3 position could produce a high level of selectivity and potency for the *P. falciparum* over the human enzyme. We also found that the adjacent P2 position could present various natural and unnatural amino acids without compromising potency. Therefore, we theorized that cyclic peptides could be synthesized via a terminal alkene on the N-terminal capping group linked through Grubbs-mediated cross metathesis to various P2 amino acids. Our first set of compounds was based on our most potent and selective linear peptide EY-4-78^11^ and contained a homophenylalanine at the P3 position with either a 2-Aminohex-5-enoic acid or an allyl protected serine at P2 to cyclize the peptides (**Figure 2A**). The linear versions of the compounds (**1**,**2**) showed equal potency against the parasite (EC_50_ < 3 nM) as compared with previously explored linear proteasome inhibitors (WLL EC_50_ = 15 nM; **Figure 2B**). The corresponding cyclized versions **(3,4)** showed similar potency to the parent molecules but the cyclization dramatically increased toxicity towards Human Foreskin Fibroblasts (HFFs) (**Supplemental Figure 1**). We altered the size of the macrocycle in the third pair of peptides by increasing the length of the N-terminal alkene, and found that changing the length of the capping group had no impact on the potency of **3** compared to linear peptides **1** and **2**. However, the 15-member macrocycle of peptide **6** had a 10-fold decrease in potency for the parasite and a 4-fold reduction in selectivity when compared to peptide **3**. This finding suggested that differences in the linear and cyclized peptides may become greater as the ring size increases (**Figure 2C**).

**Figure 2.**
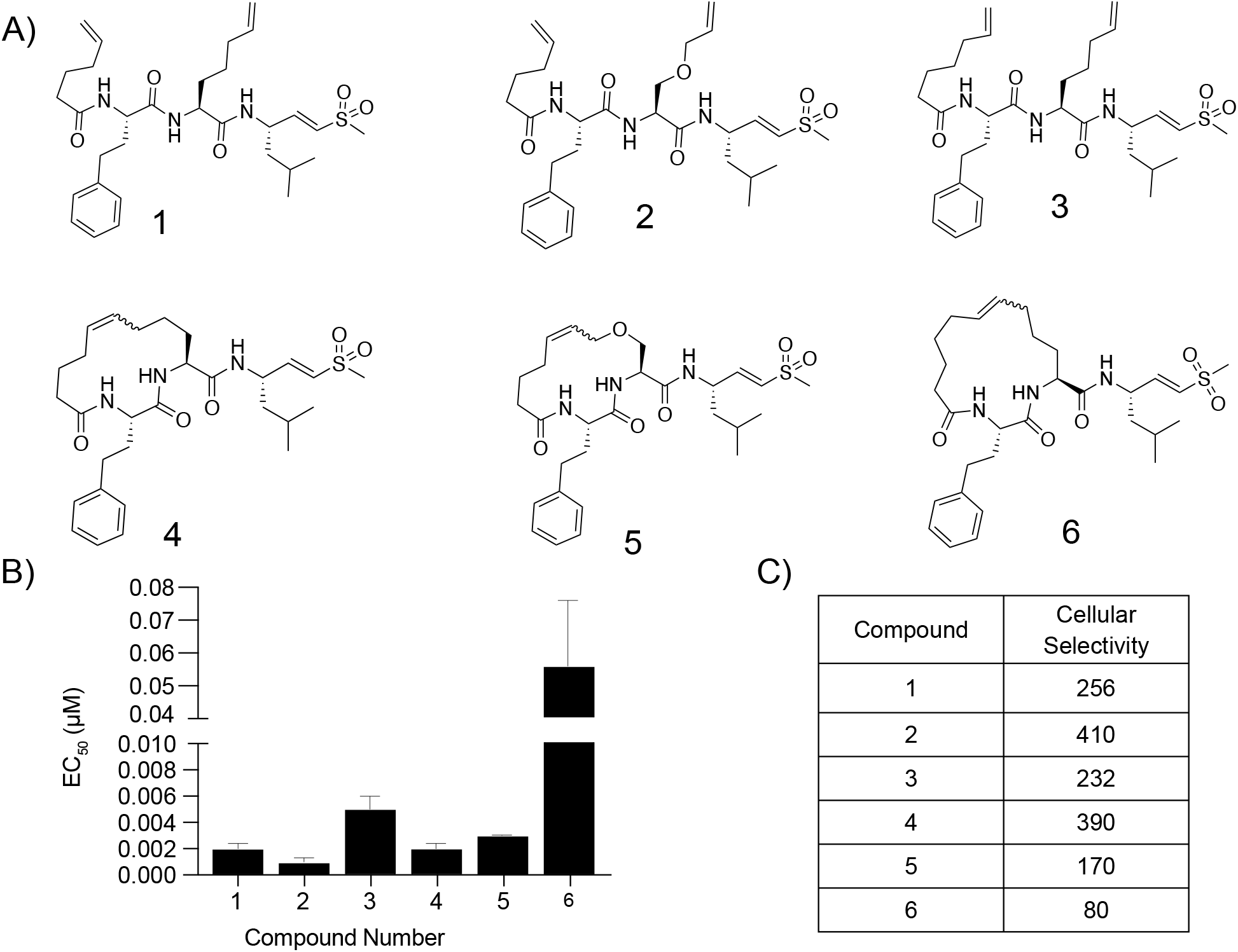
Potency and selectivity of macrocyclic peptide inhibitors containing an alkyl backbone and their linear counterparts. (A) Structures of the macrocyclic peptide inhibitors and their linear counterparts. (B) Compound mean ± SEM EC_50_ values against *P. falciparum* asexual blood stage W2 parasites of each inhibitor. Data were generated through 72 hour drug treatment assays (N, n = 2,2). (C) Table of selectivity indexes for *P. falciparum* parasites compared to primary mammalian HFF cells (listed as the ratio of HFF EC_50_ to parasite EC_50_ values).

We hypothesized that because parasitized red blood cells tend to be more permeant to peptide-based inhibitors compared to other cell types, the increased host cell toxicity in the HFFs may be due to increased lipophilicity and thus greater permeability of the cyclic peptides into human cells.^20,21^ We evaluated this hypothesis by comparing the biochemical selectivity of our compounds against purified *Plasmodium* and human proteasomes. To test the importance of lipophilicity in potency and cytotoxicity, we synthesized a second set of cyclic peptides using a diether linkage to increase the hydrophilicity of the backbone (**Figure 3A; Supplementary Scheme 1**). Interestingly, linear peptide **7** showed micromolar inhibition of parasite growth (EC_50_ = 2.7 µM) and very low HFF toxicity (EC_50_ ≥ 50 µM), while the extension of the capping group by one methylene group in peptide **8** led to both substantially more potent antiparasitic activity (EC50 = 0.016 µM), but also higher cytotoxicity (EC_50_ = 1.1 µM) (**Figure 3B and C; Supplemental Figure 2**). Peptide **8** showed modest inhibition of the *P. falciparum* proteasome β2 subunit, while peptide **7** had no inhibitory effects, and neither peptide had any inhibitory activity against the human β2 subunit (**Figure 3D**). Peptide **7** showed comparable inhibition of the β5 subunits of each proteasome and peptide **8** was 6-fold more selective for the *Plasmodium* β5 subunit over the human counterpart. Macrocycle **9** was 19-fold more potent than its linear counterpart **8** and maintained low cytotoxicity. The cyclization of **8** led to a significant decrease in cytotoxicity for macrocycle **10** while maintaining similar parasite growth inhibition, potentially due to its 3-fold decrease in potency against the purified human β5 subunit. Nevertheless, it was difficult to achieve high degrees of selectivity for purified proteasomes and it is likely that the combination of cell permeability and biochemical selectivity between the two proteasomes led to differences in antiparasitic effects and cytotoxicity. We also reduced the double bond within the macrocycles to test the importance of unsaturation within the backbone, resulting in a 2-fold decrease in potency in the saturated macrocycle **11** compared to parent compound **9**, and a 15-fold reduction in potency in compound **12** compared to the parent **10**. Both saturated macrocycles had negligible changes in cytotoxicity, suggesting that an unsaturated double bond increases the potency but not the cytotoxicity for the ether linked macrocycles.

**Figure 3.**
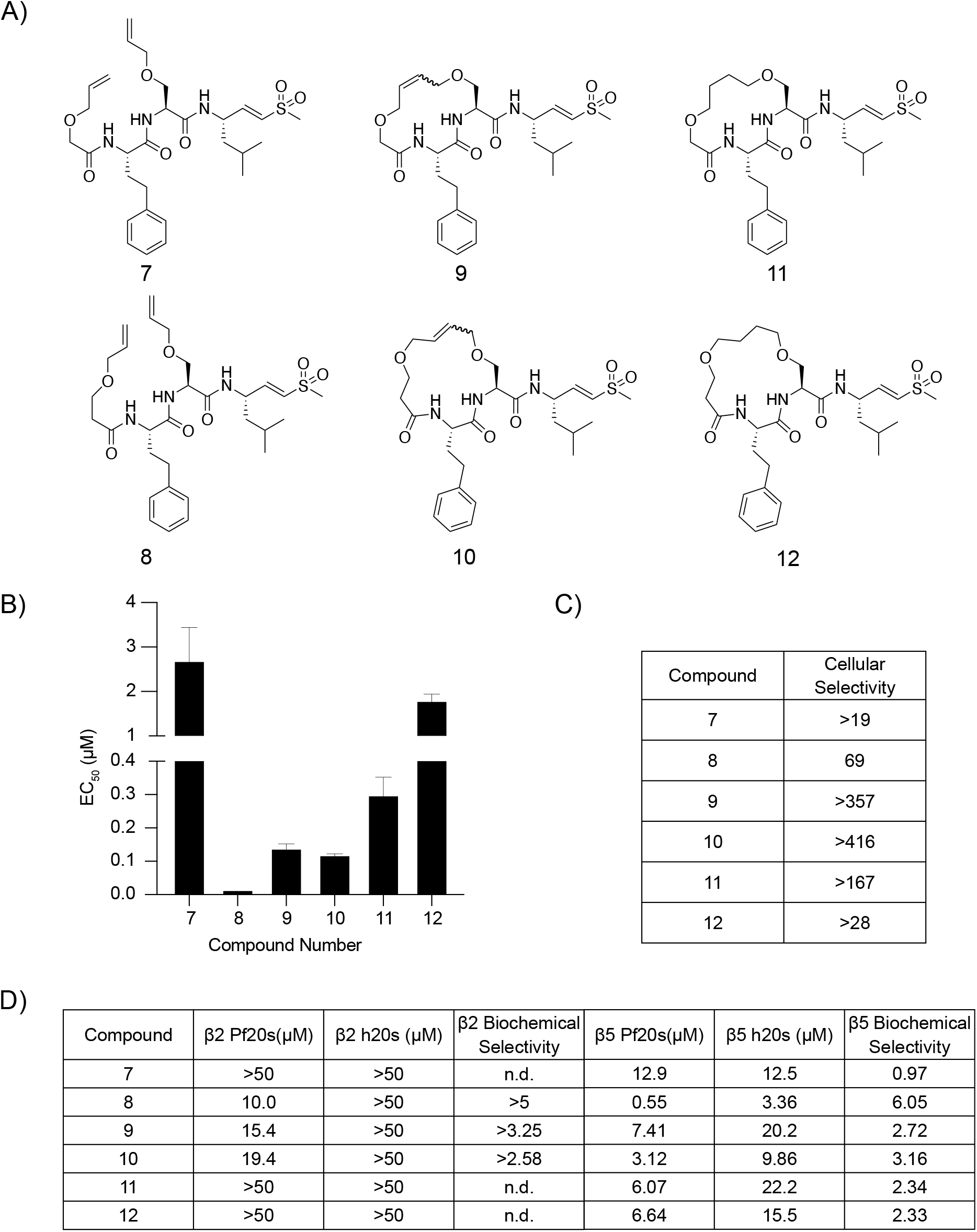
Potency and selectivity of ether linked macrocyclic peptide inhibitor and their linear counterparts. (A) Structures of ether linked cyclic peptides and corresponding linear counterparts. (B) Compound mean ± SEM EC_50_ values against *P. falciparum* asexual blood stage W2 parasites for each inhibitor. Data were generated through 72 hour drug treatment assays (N, n = 2,2). (C) Table of cellular selectivity indexes for *P. falciparum* parasites compared to primary mammalian HFF cells (listed as the ratio of HFF EC_50_ to parasite EC_50_ values). (D) Table of potencies (IC_50_) of compound against either the β2 or the β5 subunits of the purified *Plasmodium* or human 20s proteasomes in a one hour inhibition assay. Biochemical selectivity is calculated by taking a ratio of potency for the two proteasomes (h20s/Pf20s).

Although our previous set of compounds showed promise, the acyl capping group could be metabolized, similar to N-acyl amino acids.^22^ Therefore, we synthesized a third set of compounds with either imidazole or pyrazole capping groups that have previously been successfully used for orally bioavailable macrocyclic proteasome inhibitors (**Figure 4A; Supplementary Scheme 2**).^23^ Our pyrazole capped linear peptides **13** and **14** both showed greater potency and reduced off-target cytotoxicity when compared to their imidazole-containing complements (**Figure 4B and C; Supplemental Figure 3**). Both the saturated and unsaturated macrocycles followed the same trend as the linear peptides, as the pyrazole capped inhibitors had greater potency when compared to the same molecules capped with an imidazole. In each case, the imidazole capped macrocycles showed more potent inhibition of the β2 subunit of the *P. falciparum* proteasome than the corresponding pyrazole inhibitors. No compound in this set had any inhibitory effect against the β2 subunit of the human proteasome at up to 50 µM (**Figure 4D**). Notably, compounds with the larger 17-member macrocycles were the most potent inhibitors of *Plasmodium* growth (**18**, EC_50_ = 19 nM and **22**, EC_50_ = 5 nM) and were at least 200-fold more potent for the parasite over the HFFs in 72-hour dose-response assays (**Figure 4B**).

**Figure 4.**
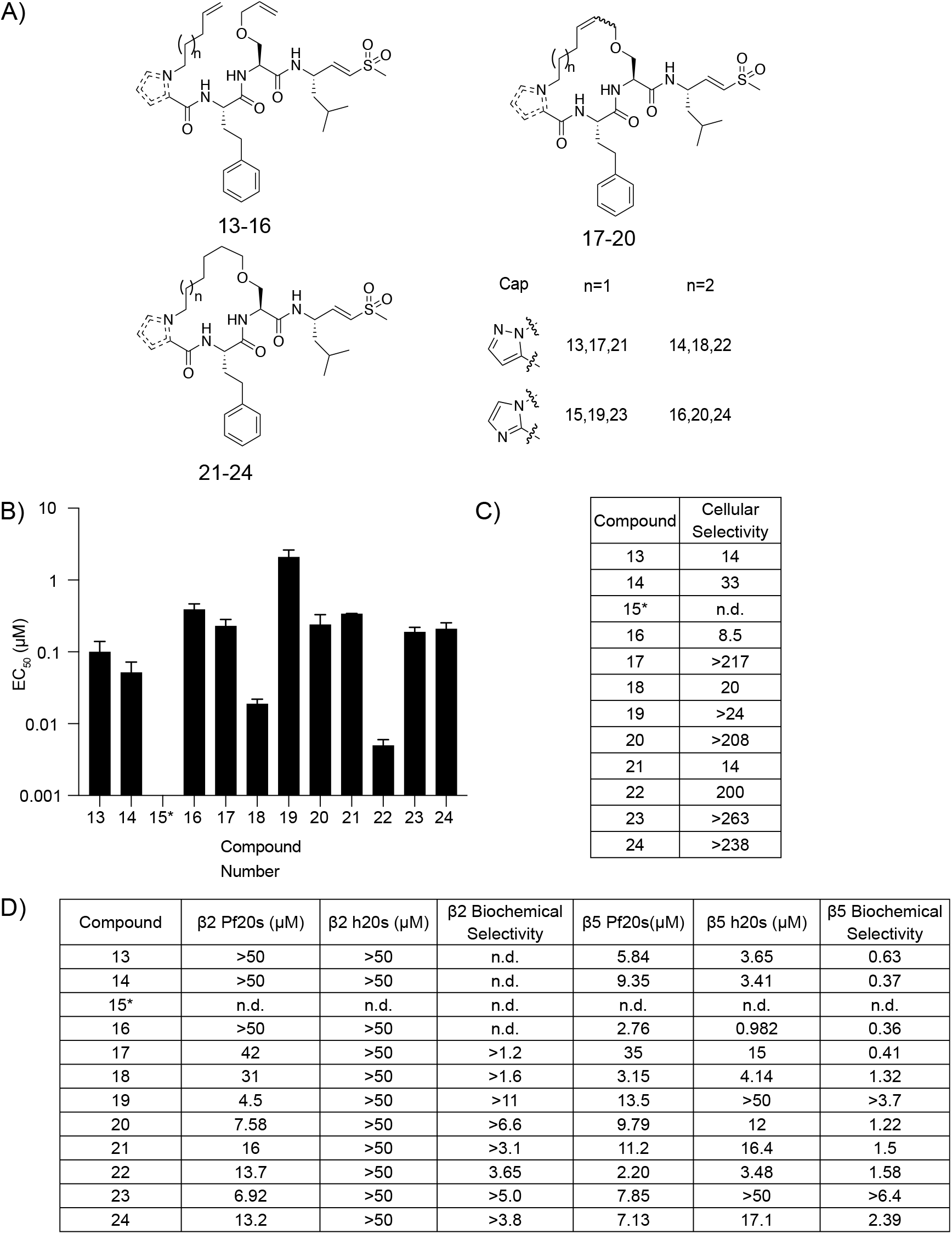
Potency and selectivity of imidazole and pyrazole capped macrocyclic peptides and their linear counterparts. (A) Structures of imidazole and pyrazole cyclic peptides and corresponding linear counterparts. The table inset shows the capping group and length of alkyl chain for each compound. (B) Compound mean ± SEM EC_50_ values against *P. falciparum* asexual blood stage W2 parasites for each inhibitor. Data were generated through 72 hour drug treatment assays (N, n = 2,2). (C) Table of cellular selectivity indexes comparing *P. falciparum* parasites to mammalian HFF cells (listed as the ratio of HFF EC_50_ to parasite EC_50_ values). (D) Table of potencies (IC_50_) of compound against either the β2 or the β5 subunits of the purified *Plasmodium* or human 20s proteasomes in a one hour inhibition assay. Biochemical selectivity is calculated by taking a ratio of potency for the two proteasomes (h20s/Pf20s).

Although **22** was slightly more cytotoxic than **18**, it was chosen to continue our structure-activity relationship study, as its synthesis does not result in isomers. Previous work had shown that the P3 homophenylalanine (hfe) is oxidized to the corresponding phenol and that the resulting phenolic metabolite has greatly reduced antiparasitic activity.^11^ To overcome this liability, we synthesized electron-deficient aromatic ring analogs of **22** to decrease the likelihood of oxidation (**Figure 5A; Supplementary Scheme 3**).^24^ Compound **26** replaced the hfe with a 4-pyridine, which resulted in a slight decrease in antiparasitic activity (EC_50_ = 75 nM), and a large improvement in cytotoxicity (EC_50_ = 20 µM) (**Figure 5B and C; Supplemental Figure 4**). Similarly, the introduction of a 4-fluorophenyl (**27**) resulted in a slight reduction in potency (EC_50_ = 39 nM) but a greater than 400-fold increase in selectivity over HFFs.

**Figure 5:**
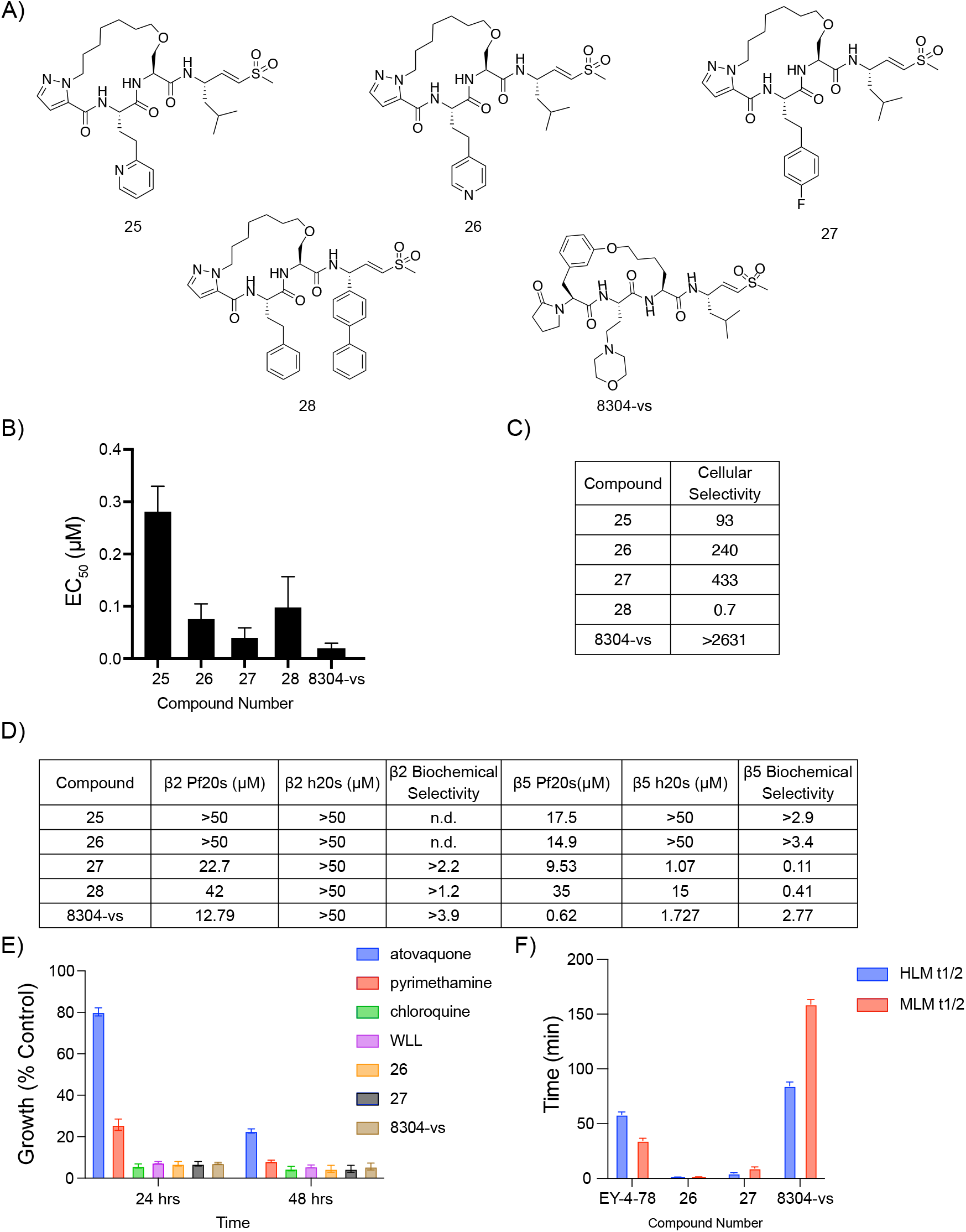
The covalent irreversible proteasome inhibitor 8304-vs is a potent, selective, and stable inhibitor of the *P. falciparum* proteasome (A) Structures of pyrazole capped inhibitors with varying P3 or P1 groups and 8304-vs, a covalent version of a previously described non-covalent macrocyclic inhibitor of the proteasome. (B) Compound mean ± SEM EC_50_ values against *P. falciparum* asexual blood stage W2 parasites for each inhibitor. Data were generated through 72 hour drug treatment assays (N, n = 2,2). (C) Table of cellular selectivity indexes comparing *P. falciparum* parasites to mammalian HFF cells (listed as the ratio of HFF EC_50_ to parasite EC_50_ values). (D) Table of potencies (IC_50_) of compound against either the β2 or the β5 subunits of the purified *Plasmodium* or human 20s proteasomes in a one-hour inhibition assay. Biochemical selectivity is calculated by taking a ratio of potency for the two proteasomes (h20s/Pf20s). (E) Plot of rate of kill assay where compounds were dosed at their EC_50_ and parasite viability was monitored over time and compared against known fast (chloroquine), medium (pyrimethamine), and slow (atovaquone) acting inhibitors of parasite growth ± SEM. N,n = 6,2. (F) Bar graphs depicting the half-life of compounds in either human (blue) or mouse (red) microsomes. Compound metabolism was measured by LC/MS/MS.

We then tested the importance of the P1 leucine moiety in potency and selectivity. Based on recent reports of potent orally available peptide boronates containing bulky P1 residues, we replaced the P1 leucine of **22** with a biphenyl alanine (**28**) (**Figure 5A; Supplementary Scheme 4**).^12^ Interestingly, this modification substantially increased the overall toxicity of the compounds towards the HFFs (**Supplemental Figure 5)**. This is likely due to the ability of the biphenyl analog to inhibit both the human β2 and β5 subunits (**Figure 5D**). We therefore did not advance this compound for further study.

To further explore the antiplasmodial effects and stability of our two most potent and selective covalent macrocyclic inhibitors (**26** and **27**) we tested them in a parasite rate of kill assay.^25^ Both inhibitors showed kill rates similar to the fast-acting antimalarial drug chloroquine, which suggests that our inhibitors could rapidly clear infections (**Figure 5E**). Parasites were treated with inhibitor for one hour, washed to remove compound and cultured a further 72 hours prior to quantifying parasite growth. Both **26** and **27** showed parasite killing at concentrations over 1 µM (**Supplemental Figure 5**). To assess metabolic stability, compounds were incubated in both human and mouse microsomes and metabolite formation was monitored over time using LC/MS. Surprisingly, **27** was metabolized quickly, with a half-life of only 1.2 minutes in both sets of microsomes. Peptide **26** was modestly better with a 4.7-minute half-life in human microsomes and 9.4 minutes in mouse microsomes (**Figure 5F**).

Given this overall poor metabolic stability of our initial lead molecules, we turned our attention to an established cyclic peptide scaffold that was highly effective at inhibiting the *Plasmodium* proteasome. This scaffold is based on a screening hit that we identified from a library of reversible inhibitors of the proteasome.^16^ The core macrocyclic scaffold was further optimized to yield a reversible inhibitor TDI-8304.^17^ We converted TDI-8304 into a covalent version by replacing the C-terminal cyclopentyl functional group with our optimal P1 leucine vinyl sulfone (**Figure 5A; Supplementary Scheme 5**). The rationale was to generate an irreversible inhibitor with increased stability compared to our original linear peptide vinyl sulfone. Furthermore, this compound allowed us to directly compare the impact of switching to a covalent inhibition mechanism on acquisition of resistance. Importantly, the resulting compound (**8304-vs**) retained the potency of the parent TDI-8304 (EC_50_ = 19 nM vs 1 nM; **Figure 5B**). In addition, the irreversible **8304-vs** showed no measurable cell toxicity within the solubility limits of the compound, suggesting that it has a > 5,000-fold selectivity for parasites over the human HFFs (**Figure 5C**). The irreversible compound also performed as well as **26** and **27** in the rate of kill assays. However, 8304-vs did not show killing at 100x EC_50_ in a one hour pulse assay (**Supplemental Figure 5**). This suggests that 8304-vs may have reduced cellular uptake or a higher dissociation constant, requiring longer incubation times to inhibit the parasite proteasome. Lastly, **8304-vs** is stable for almost 1.5 hours in human microsomes and over 2.5 hours in mouse microsomes, suggesting that it has favorable pharmacological properties and was therefore worthy of further development (**Figure 5F**).

### Resistance Selections

We profiled both **27** and **8304-vs** for their propensity to select for parasite resistance, using a minimum inoculum of resistance (MIR) platform.^26^ *P. falciparum* Dd2-B2 parasites were exposed to 3×EC_50_ concentrations of either compound at various starting inocula (four wells of 2.5×10^6^ plus three wells of 3×10^7^ parasites), and cultures were monitored for recrudescence for 60 days. No recrudescence was observed for either compound. In a repeat experiment, each compound was tested against triplicate wells of 3.3×10^7^ parasites each. One of three wells became parasite-positive for compound **27** (MIR of 1×10^8^ in this experiment), whereas no recrudescence was observed for **8304-vs**. We then performed an additional selection experiment with **8304-vs**, using two flasks of 1×10^9^ parasites each, which yielded one positive flask (MIR: 2×10^9^). An additional selection was performed with one flask of 1×10^9^ parasites from the hypermutable *P. falciparum* line, Dd2-Polδ, which also yielded recrudescent parasites. This line harbors two mutations introduced into its DNA polymerase δ and is ∼5 to 8-fold more mutable than Dd2-B2.^27^ Recrudescent parasites from selections with **27** and **8304-vs** were cloned by limiting dilution, and whole-genome sequencing of resistant clones revealed mutations in the chymotrypsin like β5 subunit and in the β6 proteasome subunit (**Table 1**). Resistance selections of **8304-vs** yielded clones with the β6 subunit mutation S157L, which was previously selected by TDI-8304 pressure.^19^ Selected clones were phenotyped and showed EC_50_ increases of 4- to 6-fold (**Supplemental Figure 6**). Compound **27** selected for the β5 subunit mutation M45I, which has previously been selected for by MPI-12.^12^ Selected clones had a 14.1- to 22.3-fold increase in their EC_50_ to **27**, compared with the parental line (**Supplemental Figure 7**).

**Table 1.** *Plasmodium falciparum* asexual blood stage minimal inoculum for resistance (MIR) selection summary.

| Name | Chemical class | Starting selective pressure | Resistant parasites in 4 wells of $2.5 \times 10^6$ | Resistant parasites in 3 wells of $3 \times 10^7$ | Resistant parasites in 2 flasks of $1 \times 10^9$ | Minimum Inoculum of Resistance (MIR) | Selected proteasome mutation | EC<sub>50</sub> fold shift against selection compound<sup>c</sup> |
| --- | --- | --- | --- | --- | --- | --- | --- | --- |
| WLL | Vinyl sulfone | 3×EC<sub>50</sub> (50 nM) | 0/4 wells | None | ND<sup>a</sup> | $>1 \times 10^9$ | N/A | NA |
| 27 | Vinyl sulfone | 3×EC<sub>50</sub> (23 nM) | 0/4 wells | 1/3 wells | ND | $1 \times 10^8$ | β5 M45I | 14.1-22.3× |
| 8304-vs | Vinyl sulfone | 3×EC<sub>50</sub> (72 nM) | 0/4 wells | 0/3 wells | 1/2 flasks<sup>b</sup> | $2 \times 10^9$ | β6 S157L | 4.0-6.0× |
| TDI-8304 | Macrocyclic peptide | 3×EC<sub>50</sub> (33 nM) | 0/4 wells | 3/3 wells | ND | $3 \times 10^7$ | β6 S157L<br>β6 N151Y | 2,621×<br>1.7× |

<sup>a</sup>Prior studies with WLL selections obtained 0/3 recrudescence flasks inoculated with $3.3 \times 10^8$ each, yielding a predicted MIR $>1 \times 10^9$ .
<sup>b</sup>One flask of the hypermutable Dd2-B2 Polδ line inoculated at $1 \times 10^9$ parasites and pressured with 8304-vs also yielded positive parasites, as reported herein.
<sup>c</sup>EC<sub>50</sub> fold shift calculated as a ratio between the EC<sub>50</sub> of the selected mutant over the EC<sub>50</sub> of the Dd2-B2 parental line. N,n = 4–5,2. ND, not done. N/A, not applicable. MIRs are calculated as the total number of parasites inoculated divided by the total number of positive cultures (when obtained). The formula extends up to the largest inoculum at which parasites were obtained (or tested in the case of negative cultures) and includes lower inocula (see Methods).

### Mutant Panel

Because the addition of the vinyl sulfone to TDI-8304 greatly improved its resistance profile, we synthesized a 2,4-diflourophenylalanine epoxyketone to assess the importance of specific covalent warheads for overcoming resistance. We profiled **8304-vs**, **8304-epoxy**, and **27** against a panel of proteasome mutants selected for in previous studies,^19^ comprising several β5 and β6 mutations on a Dd2-B2 background. Compound **27** had a significant increase in EC_50_ against all proteasome mutant lines assessed, with the exception of β5 A50V and β6 N151Y (the latter of which saw a slight but significant decrease in EC_50_ (**Supplemental Figure 8**). Overall, **8304-vs** showed a modest increase in EC_50_ in parasites with the β5 M45I and M45V mutations (1.8 fold and 1.5 fold respectively), and a slightly higher 8.8-fold increase in EC_50_ in parasites with the β6 S157L mutation (**Figure 6**). Interestingly, this mutation was also present in previous selections with TDI-8304, to which it mediated a more than 2,900-fold increase in EC_50._^19^ None of the mutants profiled had significantly increased EC_50_ values to 8304-epoxy, and a slight decrease in the EC_50_ value for this compound was observed with β5 M45V mutant parasites.

**Figure 6.**
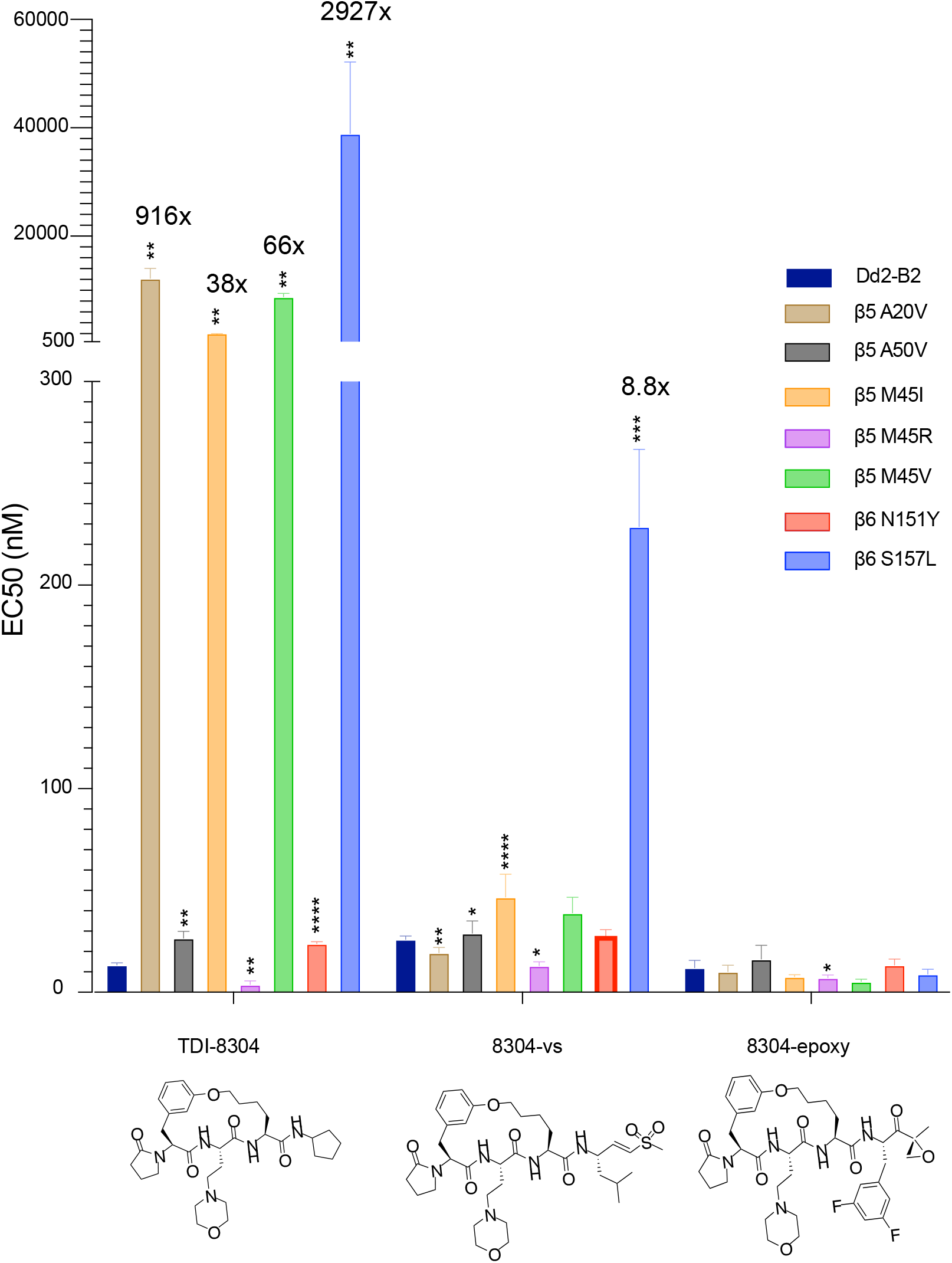
Profiling of potency of macrocyclic reversible and irreversible proteasome inhibitor based on the TDI-8304 scaffold against *P. falciparum* 20S β5 and β6 subunit mutants. Dd2-B2 parasites containing previously selected proteasome mutations were assayed against TDI-8304 (PMID 36963402) and the irreversible analogs 8304-vs and 8304-Eps. Statistical significance, compared to Dd2-B2, was calculated using unpaired t tests with Welch’s correction. *p<0.05; **p<0.01; ***p<0.001; ****p<0.0001. N,n = 3-5,2.

## DISCUSSION

Resistance to ART and ACT treatments is a major challenge to the control and mitigation of malaria. The *Plasmodium* proteasome has emerged as a high-value drug target due to its necessity for parasite development and lifecycle progression, and for the essential role it plays in reducing ART-induced proteotoxic stress. Proteasome inhibitors also have the very desirable feature of synergizing with ART derivatives, including against ART-resistant parasites.^9,13^ Combination therapies utilizing proteasome inhibitors in tandem with ART are, therefore, a promising strategy both to treat malaria and combat the dissemination of ART resistance. Nevertheless, the challenge remains to design *Plasmodium* proteasome-specific inhibitors that do not cross react with human proteasomes. Previous work in our lab leveraged the use of natural and unnatural amino acids to design selective irreversible covalent *P. falciparum* inhibitors.^9,11^ Having previously defined the P1 and P3 peptide positions as the determinants of selectivity, we describe here the synthesis and characterization of irreversible covalent macrocycles linked via the P2 side chain and the N-terminal capping group. We were able to determine the impact of various capping groups and several types of cycle linkages on antiparasitic potency and off-target mammalian cytotoxicity. Notably, decreasing the lipophilicity of our cycles resulted in reduced cytotoxicity. We also found that substituting the capping group of our cycles with a pyrazole greatly increased the potency towards the parasite, but also increased cytotoxicity. Interestingly, exchange of our P1 leucine with a biphenyl alanine resulted in a complete inversion of selectivity, both phenotypically and biochemically, which underscores the importance of the P1 leucine for *Plasmodium* selectivity. This observation suggests that moieties similar to the bulky biphenyl alanine vinyl sulfone at the P1 position could be used to develop better inhibitors of the human proteasome. We sought to optimize our inhibitors for metabolic stability by varying electron-deficient aromatic side chains on the P3. While these analogs proved to be potent inhibitors of parasite growth with low cytotoxicity, they ultimately showed poor microsomal stability, suggesting that cyclization alone cannot be used to increase metabolic stability.

We subsequently focused our attention on developing an irreversible covalent inhibitor, based on a structure previously explored for non-covalent inhibition of the *Plasmodium* proteasome.^17^ This strategy has proven to be powerful in developing covalent kinase inhibitors by appending an electrophilic reactive group to non-covalent compounds.^28^ We posited that adding a vinyl sulfone onto the P1 position of the cycle could allow it to react with the active site threonine and convert the reversible inhibitor TDI-8304 into the covalent irreversible inhibitor 8304-vs. The resulting compound showed comparable potency to TDI-8304, with no cytotoxicity to HFFs, a fast rate of kill, and substantially more stability than previous linear peptide inhibitors and the cyclic peptides tested herein.

Converting the previously tested non-covalent proteasome inhibitor TDI-8304 into the irreversible covalent inhibitor 8304-vs, via the addition of the vinyl sulfone reactive group, enabled us to compare their resistance liabilities as a function of the inhibition mechanism. The irreversible 8304-vs showed a MIR of 2×10^9^, compared to our earlier value of 3×10^7^ obtained with TDI-8304.^19^ This new vinyl sulfone therefore yielded a very similar resistance profile to WLL-vs, for which resistance was only obtained with 2×10^9^ parasites.^13,19^ These data provide evidence for a significantly improved resistance liability associated with covalent binding. For the reversible inhibitor TDI-8304, mutations in the proteasome likely increase the off rate of the compound, leading to reduced active site occupancy and lower overall inhibition as the molecule is cleared. While the off rate for the irreversibly binder 8304-vs can also be increased by mutations within the proteasome, covalent inhibitors work via a two-step inhibitory reaction - a reversible association followed by an irreversible covalent bond formation.^29^ The covalent 8304-vs may therefore experience similar off rates to its non-covalent counterpoint, but over time, it will irreversibly inactivate the proteasome. This is clearly demonstrated with the S157L mutation, which was selected for by both TDI-8304 and 8304-vs, yet showed a 2,900-fold increase in EC_50_ for the former, contrasting with a 4- to 6-fold shift for the latter. This mutation likely leads to the loss of a hydrogen bond donor and acceptor from the serine hydroxyl group required for TDI-8304 binding to the proteasome active site, and 8304-vs is presumably less impacted by this mutation because of its electrophilic warhead. 8304-vs also compared favorably to TDI-8304 against a panel of parasite lines with mutations in the *P. falciparum* proteasome subunits β5 and β6, showing increases in EC_50_ that were orders of magnitude lower than the fold shifts seen against TDI-8304.

To further demonstrate the importance of covalency to overcome resistance we synthesized an epoxyketone derivative, 8304-epoxy. This compound performed extremely well against the panel of proteasome mutant lines, with no significant increase in EC_50_ against any of the lines tested. However, it was much less stable than 8304-vs in both human and mouse microsomes (**Supplemental Figure 9**). The instability of 8304-epoxy is likely due to the susceptibility of the epoxy ring opening as well as the epoxyketone being a substrate for other enzymes such as serine hydrolases, while the vinyl sulfone is much more stable due to its lower general reactivity.^30,31^ These results reinforce our previous work in suggesting that the vinyl sulfone is the optimal electrophilic warhead for the development of an irreversible covalent proteasome drug inhibitor.

## CONCLUSIONS

Novel treatments and strategies are needed to combat the growth of ART-resistant parasites. The *P. falciparum* proteasome, an enzyme essential at all parts of the parasite’s life cycle, continues to be a promising target for such therapies. Our data provide compelling evidence that optimized properties for both covalent and noncovalent macrocyclic inhibitors of the *Plasmodium* proteasome can be combined to create more potent and selective inhibitors of parasite growth with better stability. Moreover, our efforts reinforce the importance of covalency in overcoming parasite resistance, and specifically the difficulty in developing resistance against covalent inhibitors of the proteasome. While **8304-vs** stands out as a lead compound, further testing is needed to determine bioavailability and *in vivo* efficacy. This underscores the importance for further synthetic and medicinal chemistry efforts to advance proteasome inhibitors as promising new drugs to partner with ART derivatives and leverage their synergy against ART-resistant parasites.

## Ancillary Information

### Supporting Information

This includes HFF Cell Viability for Compounds (Figures S1-S4), Pulse Killing Assays (Figure S5), Minimum Inoculum of Resistance selections (Figures S6-S8), microsome stability (Figure S9), compound characterization and compound synthesis (Supplementary Schemes), and Mass Spectrometry analysis.

## Acknowledgment

This work was supported in part by the NIH (R21/R33 AI125781 to M.B. and D.A.F.; R01 AI109023 to D.A.F.; and R01 AI143714 to G.L.). Additional funding was provided by the Department of Defense (W81XWH2210520 to D.A.F., M.B., and W.H.G.).

## Methods

### Mammalian Cytotoxicity Assays

Human Foreskin Fibroblasts (HFFs) were kept in DMEM supplemented with 5% FBS. HFFs were seeded at 2000 cells per well for 4 hours and then compound was added for 72 hours. Cell viability was measured using the CellTiter-Blue Assay (Promega) as per manufacturer’s instructions.

### Biochemical Inhibition of the Proteasome

Purified Pf20S or h20S proteasomes were both tested for subunit specific inhibition in 20 mM HEPES and 0.5 mM EDTA. For β2 inhibition, purified proteasome (1 nM final concentration) was activated with PA28 (12nM final concentration; E380 Boston Biochem) followed by addition of Boc-LRR-AMC (50 µM final concentration) and inhibitor simultaneously Cleavage was measured using fluorescence (EX 380 nm/EM 460 nm) with Cytation 3 imaging reader (BioTek, Winooski, VT, USA) for 60 minutes and slope was used to calculate the inhibition curve. For β5 inhibition, the Suc-LLVY-AMC (10 µM final concentration) was used as the substrate.

### *In vitro* drug susceptibility assays

In the Bogyo lab, *P. falciparum* W2 parasites, obtained from the Malaria Research and Reference Reagent Resource Center, were cultured in human erythrocytes purchased from the Stanford Blood Center (No blood type discrimination) and maintained as previously described.^7^ Ring stage *P. falciparum* were inoculated at 1% parasitemia and 0.5% hematocrit in a 96-well plate spotted with compounds. The cultures were incubated in a 10-point two-fold serial dilution dose-response for 72 hours and then fixed with a final concentration of 1% paraformaldehyde (in PBS) for 30 minutes at room temperature. The nuclei were stained with YOYO-1 at a final concentration of 50 nM and incubated at room temperature overnight. The percentage of YOYO-1 positive parasitized erythrocytes was quantified per well by a BD Accuri C6 automated flow cytometer. In pulse assays, plates were incubated with compound for 1 hour. Parasites were then washed twice with media, before fresh media was added, and growth continued for 71 hours prior to quantifying parasitemia as above. Fosmidomycin was used as a positive control. In the Fidock Lab, the sensitivity of Dd2-B2 proteasome WT and mutant lines to our compounds was determined by exposing asexual blood stage parasites to a 10-point two-fold serial dilution dose-response assay. Parasites were seeded at 0.2% parasitemia and 1% hematocrit, and plated in 96-well plates with a final volume of 200 µL. Plates were incubated at 37°C for 72 hours under normal culturing conditions. Final parasitemia was determined through flow cytometry on an iQue Plus flow cytometer following staining with 1×SYBR Green and 100 nM MitoTracker Deep Red (ThermoFisher) for 30 minutes at 37°C. Statistical significance was calculated through unpaired t tests with Welch’s correction.

### Parasite culture for resistance

*P. falciparum* asexual blood stage parasites were cultured at 3% hematocrit in human O+ RBCs in RPMI-1640 media, supplemented with 25 mM HEPES, 50 mg/L-hypoxanthine, 2mM L-glutamine, 0.21% sodium bicarbonate, 0.5% (wt/vol) AlbuMAXII (Invitrogen) and 10 μg/mL gentamycin, in modular incubator chambers (Billups-Rothenberg) at 5% O2, 5% CO2 and 90% N2 at 37°C. Dd2 parasites were obtained from T. Wellems (NIAID, NIH). Dd2-B2 is a genetically homogenous line that was cloned from Dd2 by limiting dilution in the Fidock lab.

### Minimum Inoculum of Resistance Studies

Resistance selections were performed by culturing Dd2-B2 or Dd2 polο parasites at 3 x EC_50_, as previously described.^19^ Media was changed daily for the first 6 days of selections, and cultures were monitored by Giemsa staining and microscopy until parasites were cleared. Thereafter, media was replenished every two days, and cultures were monitored twice weekly for recrudescence by Giemsa staining and microscopy. Cultures were maintained under drug pressure for 60 days or until recrudescent parasites were observed. Bulk resistant lines were cloned by limited dilution cloning. The MIR value is defined as the minimum number of parasites used to obtain resistance and calculated as follows: total number of parasites inoculated ÷ total number of positive cultures.

### Whole Genome Sequencing

*P. falciparum* parasites were lysed in 0.05% saponin and washed with 1×PBS, and genomic DNA (gDNA) was purified using the QIAamp DNA Blood Midi Kit (Qiagen). gDNA concentrations were quantified by Qubit using the dsDNA HS Assay (Invitrogen). 200ng of gDNA was used to prepare sequencing libraries using the Illumina DNA Prep kit with Nextera™ DNA CD Indexes (Illumina). Samples were multiplexed and sequenced on an Illumina MiSeq using the MiSeq Reagent Kit V3 600 (Illumina) to obtain 300 base pair paired-end reads at an average of 30× depth of coverage. Sequence reads were aligned to the *P. falciparum* 3D7 reference genome (PlasmoDB version 48) using Burrow-Wheeler Alignment. PCR duplicates and unmapped reads were filtered out using Samtools and Picard. Reads were realigned around indels using GATK RealignerTargetCreator, and base quality scores were recalibrated using GATK BaseRecalibrator. GATK HaplotypeCaller (version 4.2.2) was used to identify all single nucleotide polymorphisms (SNPs). SNPs were filtered based on quality scores (variant quality as function of depth QD >1.5, mapping quality >40, min base quality score >18) and read depth (>5) to obtain high-quality SNPs, which were annotated using snpEFF. Integrated Genome Viewer was used to visually verify the presence of SNPs. BIC-Seq was used to check for copy number variations using the Bayesian statistical model. Copy number variations in highly polymorphic surface antigens and multi-gene families were removed as these are prone to stochastic changes during *in vitro* culture.

## References

(1) World Health Organization. World Malaria Report 2022. 2022. https://www.who.int/teams/global-malaria-programme/reports/world-malaria-report-2022

(2) Noedl, H.; Socheat, D.; Satimai, W. Artemisinin-resistant malaria in Asia. N. Engl. J. Med. 2009, 361 (5), 540–541. https://doi.org/10.1056/NEJMc0900231.

(3) Uwimana, A.; Legrand, E.; Stokes, B. H.; Ndikumana, J.-L. M.; Warsame, M.; Umulisa, N.; Ngamije, D.; Munyaneza, T.; Mazarati, J.-B.; Munguti, K.; Campagne, P.; Criscuolo, A.; Ariey, F.; Murindahabi, M.; Ringwald, P.; Fidock, D. A.; Mbituyumuremyi, A.; Menard, D. Emergence and Clonal Expansion of *in Vitro* Artemisinin-Resistant Plasmodium Falciparum Kelch13 R561H Mutant Parasites in Rwanda. Nat. Med. 2020, 26 (10), 1602–1608. https://doi.org/10.1038/s41591-020-1005-2.

(4) Balikagala, B.; Fukuda, N.; Ikeda, M.; Katuro, O. T.; Tachibana, S.-I.; Yamauchi, M.; Opio, W.; Emoto, S.; Anywar, D. A.; Kimura, E.; Palacpac, N. M. Q.; Odongo-Aginya, E. I.; Ogwang, M.; Horii, T.; Mita, T. Evidence of Artemisinin-Resistant Malaria in Africa. N. Engl. J. Med. 2021, 385 (13), 1163–1171. https://doi.org/10.1056/NEJMoa2101746.

(5) Straimer, J.; Gnädig, N. F.; Witkowski, B.; Amaratunga, C.; Duru, V.; Ramadani, A. P.; Dacheux, M.; Khim, N.; Zhang, L.; Lam, S.; Gregory, P. D.; Urnov, F. D.; Mercereau-Puijalon, O.; Benoit-Vical, F.; Fairhurst, R. M.; Ménard, D.; Fidock, D. A. K13-Propeller Mutations Confer Artemisinin Resistance in *Plasmodium Falciparum* Clinical Isolates. Science 2015, 347 (6220), 428–431. https://doi.org/10.1126/science.1260867.

(6) Ashley, E. A.; Dhorda, M.; Fairhurst, R. M.; Amaratunga, C.; Lim, P.; Suon, S.; Sreng, S.; Anderson, J. M.; Mao, S.; Sam, B.; Sopha, C.; Chuor, C. M.; Nguon, C.; Sovannaroth, S.; Pukrittayakamee, S.; Jittamala, P.; Chotivanich, K.; Chutasmit, K.; Suchatsoonthorn, C.; Runcharoen, R.; Hien, T. T.; Thuy-Nhien, N. T.; Thanh, N. V.; Phu, N. H.; Htut, Y.; Han, K.-T.; Aye, K. H.; Mokuolu, O. A.; Olaosebikan, R. R.; Folaranmi, O. O.; Mayxay, M.; Khanthavong, M.; Hongvanthong, B.; Newton, P. N.; Onyamboko, M. A.; Fanello, C. I.; Tshefu, A. K.; Mishra, N.; Valecha, N.; Phyo, A. P.; Nosten, F.; Yi, P.; Tripura, R.; Borrmann, S.; Bashraheil, M.; Peshu, J.; Faiz, M. A.; Ghose, A.; Hossain, M. A.; Samad, R.; Rahman, M. R.; Hasan, M. M.; Islam, A.; Miotto, O.; Amato, R.; MacInnis, B.; Stalker, J.; Kwiatkowski, D. P.; Bozdech, Z.; Jeeyapant, A.; Cheah, P. Y.; Sakulthaew, T.; Chalk, J.; Intharabut, B.; Silamut, K.; Lee, S. J.; Vihokhern, B.; Kunasol, C.; Imwong, M.; Tarning, J.; Taylor, W. J.; Yeung, S.; Woodrow, C. J.; Flegg, J. A.; Das, D.; Smith, J.; Venkatesan, M.; Plowe, C. V.; Stepniewska, K.; Guerin, P. J.; Dondorp, A. M.; Day, N. P.; White, N. J. Spread of Artemisinin Resistance in *Plasmodium Falciparum* Malaria. N. Engl. J. Med. 2014, 371 (5), 411–423. https://doi.org/10.1056/NEJMoa1314981.

(7) Li, H.; Ponder, E. L.; Verdoes, M.; Asbjornsdottir, K. H.; Deu, E.; Edgington, L. E.; Lee, J. T.; Kirk, C. J.; Demo, S. D.; Williamson, K. C.; Bogyo, M. Validation of the Proteasome as a Therapeutic Target in *Plasmodium* Using an Epoxyketone Inhibitor with Parasite-Specific Toxicity. Chem. Biol. 2012, 19 (12), 1535–1545. https://doi.org/10.1016/j.chembiol.2012.09.019.

(8) Bridgford, J. L.; Xie, S. C.; Cobbold, S. A.; Pasaje, C. F. A.; Herrmann, S.; Yang, T.; Gillett, D. L.; Dick, L. R.; Ralph, S. A.; Dogovski, C.; Spillman, N. J.; Tilley, L. Artemisinin Kills Malaria Parasites by Damaging Proteins and Inhibiting the Proteasome. Nat. Commun. 2018, 9 (1), 3801. https://doi.org/10.1038/s41467-018-06221-1.

(9) Li, H.; O’Donoghue, A. J.; van der Linden, W. A.; Xie, S. C.; Yoo, E.; Foe, I. T.; Tilley, L.; Craik, C. S.; da Fonseca, P. C. A.; Bogyo, M. Structure- and Function-Based Design of *Plasmodium*-Selective Proteasome Inhibitors. Nature 2016, 530 (7589), 233–236. https://doi.org/10.1038/nature16936.

(10) LaMonte, G. M.; Almaliti, J.; Bibo-Verdugo, B.; Keller, L.; Zou, B. Y.; Yang, J.; Antonova-Koch, Y.; Orjuela-Sanchez, P.; Boyle, C. A.; Vigil, E.; Wang, L.; Goldgof, G. M.; Gerwick, L.; O’Donoghue, A. J.; Winzeler, E. A.; Gerwick, W. H.; Ottilie, S. Development of a Potent Inhibitor of the *Plasmodium* Proteasome with Reduced Mammalian Toxicity. J. Med. Chem. 2017, 60 (15), 6721–6732. https://doi.org/10.1021/acs.jmedchem.7b00671.

(11) Yoo, E.; Stokes, B. H.; de Jong, H.; Vanaerschot, M.; Kumar, T.; Lawrence, N.; Njoroge, M.; Garcia, A.; Van der Westhuyzen, R.; Momper, J. D.; Ng, C. L.; Fidock, D. A.; Bogyo, M. Defining the Determinants of Specificity of *Plasmodium* Proteasome Inhibitors. J. Am. Chem. Soc. 2018, 140 (36), 11424–11437. https://doi.org/10.1021/jacs.8b06656.

(12) Xie, S. C.; Metcalfe, R. D.; Mizutani, H.; Puhalovich, T.; Hanssen, E.; Morton, C. J.; Du, Y.; Dogovski, C.; Huang, S.-C.; Ciavarri, J.; Hales, P.; Griffin, R. J.; Cohen, L. H.; Chuang, B.-C.; Wittlin, S.; Deni, I.; Yeo, T.; Ward, K. E.; Barry, D. C.; Liu, B.; Gillett, D. L.; Crespo-Fernandez, B. F.; Ottilie, S.; Mittal, N.; Churchyard, A.; Ferguson, D.; Aguiar, A. C. C.; Guido, R. V. C.; Baum, J.; Hanson, K. K.; Winzeler, E. A.; Gamo, F.-J.; Fidock, D. A.; Baud, D.; Parker, M. W.; Brand, S.; Dick, L. R.; Griffin, M. D. W.; Gould, A. E.; Tilley, L. Design of Proteasome Inhibitors with Oral Efficacy *in Vivo* against *Plasmodium Falciparum* and Selectivity over the Human Proteasome. Proc. Natl. Acad. Sci. 2021, 118 (39), e2107213118. https://doi.org/10.1073/pnas.2107213118.

(13) Stokes, B. H.; Yoo, E.; Murithi, J. M.; Luth, M. R.; Afanasyev, P.; Da Fonseca, P. C. A.; Winzeler, E. A.; Ng, C. L.; Bogyo, M.; Fidock, D. A. Covalent *Plasmodium falciparum*-Selective Proteasome Inhibitors Exhibit a Low Propensity for Generating Resistance *in Vitro* and Synergize with Multiple Antimalarial Agents. PLOS Pathog. 2019, 15 (6), e1007722. https://doi.org/10.1371/journal.ppat.1007722.

(14) Lau, J. L.; Dunn, M. K. Therapeutic Peptides: Historical Perspectives, Current Development Trends, and Future Directions. Bioorg. Med. Chem. 2018, 26 (10), 2700–2707. https://doi.org/10.1016/j.bmc.2017.06.052.

(15) Nielsen, D. S.; Shepherd, N. E.; Xu, W.; Lucke, A. J.; Stoermer, M. J.; Fairlie, D. P. Orally Absorbed Cyclic Peptides. Chem. Rev. 2017, 117 (12), 8094–8128. https://doi.org/10.1021/acs.chemrev.6b00838.

(16) Li, H.; Tsu, C.; Blackburn, C.; Li, G.; Hales, P.; Dick, L.; Bogyo, M. Identification of Potent and Selective Non-Covalent Inhibitors of the *Plasmodium falciparum* Proteasome. J. Am. Chem. Soc. 2014, 136 (39), 13562–13565. https://doi.org/10.1021/ja507692y.

(17) Zhan, W.; Zhang, H.; Ginn, J.; Leung, A.; Liu, Y. J.; Michino, M.; Toita, A.; Okamoto, R.; Wong, T.; Imaeda, T.; Hara, R.; Yukawa, T.; Chelebieva, S.; Tumwebaze, P. K.; Lafuente-Monasterio, M. J.; Martinez-Martinez, M. S.; Vendome, J.; Beuming, T.; Sato, K.; Aso, K.; Rosenthal, P. J.; Cooper, R. A.; Meinke, P. T.; Nathan, C. F.; Kirkman, L. A.; Lin, G. Development of a Highly Selective *Plasmodium falciparum* Proteasome Inhibitor with Anti-malaria Activity in Humanized Mice. Angew. Chem. Int. Ed. 2021, 60 (17), 9279–9283. https://doi.org/10.1002/anie.202015845.

(18) Zhang, H.; Ginn, J.; Zhan, W.; Liu, Y. J.; Leung, A.; Toita, A.; Okamoto, R.; Wong, T.-T.; Imaeda, T.; Hara, R.; Yukawa, T.; Michino, M.; Vendome, J.; Beuming, T.; Sato, K.; Aso, K.; Meinke, P. T.; Nathan, C. F.; Kirkman, L. A.; Lin, G. Design, Synthesis, and Optimization of Macrocyclic Peptides as Species-Selective Antimalaria Proteasome Inhibitors. J. Med. Chem. 2022, 65 (13), 9350–9375. https://doi.org/10.1021/acs.jmedchem.2c00611.

(19) Deni, I.; Stokes, B. H.; Ward, K. E.; Fairhurst, K. J.; Pasaje, C. F. A.; Yeo, T.; Akbar, S.; Park, H.; Muir, R.; Bick, D. S.; Zhan, W.; Zhang, H.; Liu, Y. J.; Ng, C. L.; Kirkman, L. A.; Almaliti, J.; Gould, A. E.; Duffey, M.; O’Donoghue, A. J.; Uhlemann, A.-C.; Niles, J. C.; Da Fonseca, P. C. A.; Gerwick, W. H.; Lin, G.; Bogyo, M.; Fidock, D. A. Mitigating the Risk of Antimalarial Resistance via Covalent Dual-Subunit Inhibition of the *Plasmodium* Proteasome. Cell Chem. Biol. 2023, S2451945623000612. https://doi.org/10.1016/j.chembiol.2023.03.002.

(20) Desai, S. A. Why Do Malaria Parasites Increase Host Erythrocyte Permeability? Trends Parasitol. 2014, 30 (3), 151–159. https://doi.org/10.1016/j.pt.2014.01.003.

(21) Naccache, P.; Sha’afi, R. I. Patterns of Nonelectrolyte Permeability in Human Red Blood Cell Membrane. J. Gen. Physiol. 1973, 62 (6), 714–736. https://doi.org/10.1085/jgp.62.6.714.

(22) Battista, N.; Bari, M.; Bisogno, T. N-Acyl Amino Acids: Metabolism, Molecular Targets, and Role in Biological Processes. Biomolecules 2019, 9 (12), 822. https://doi.org/10.3390/biom9120822.

(23) Li, D.; Zhang, X.; Ma, X.; Xu, L.; Yu, J.; Gao, L.; Hu, X.; Zhang, J.; Dong, X.; Li, J.; Liu, T.; Zhou, Y.; Hu, Y. Development of Macrocyclic Peptides Containing Epoxyketone with Oral Availability as Proteasome Inhibitors. J. Med. Chem. 2018, 61 (20), 9177–9204. https://doi.org/10.1021/acs.jmedchem.8b00819.

(24) Lazzara, P. R.; Moore, T. W. Scaffold-Hopping as a Strategy to Address Metabolic Liabilities of Aromatic Compounds. RSC Med. Chem. 2020, 11 (1), 18–29. https://doi.org/10.1039/C9MD00396G.

(25) Sanz, L. M.; Crespo, B.; De-Cózar, C.; Ding, X. C.; Llergo, J. L.; Burrows, J. N.; García-Bustos, J. F.; Gamo, F.-J. *P. falciparum In Vitro* Killing Rates Allow to Discriminate between Different Antimalarial Mode-of-Action. PLoS ONE 2012, 7 (2), e30949. https://doi.org/10.1371/journal.pone.0030949.

(26) Duffey, M.; Blasco, B.; Burrows, J. N.; Wells, T. N. C.; Fidock, D. A.; Leroy, D. Assessing Risks of *Plasmodium falciparum* Resistance to Select Next-Generation Antimalarials. Trends Parasitol. 2021, 37 (8), 709–721. https://doi.org/10.1016/j.pt.2021.04.006.

(27) Kümpornsin, K.; Kochakarn, T.; Yeo, T.; Okombo, J.; Luth, M. R.; Hoshizaki, J.; Rawat, M.; Pearson, R. D.; Schindler, K. A.; Mok, S.; Park, H.; Uhlemann, A.-C.; Jana, G. P.; Maity, B. C.; Laleu, B.; Chenu, E.; Duffy, J.; Moliner Cubel, S.; Franco, V.; Gomez-Lorenzo, M. G.; Gamo, F. J.; Winzeler, E. A.; Fidock, D. A.; Chookajorn, T.; Lee, M. C. S. Generation of a Mutator Parasite to Drive Resistome Discovery in *Plasmodium falciparum*. Nat. Commun. 2023, 14 (1), 3059. https://doi.org/10.1038/s41467-023-38774-1.

(28) Abdeldayem, A.; Raouf, Y. S.; Constantinescu, S. N.; Moriggl, R.; Gunning, P. T. Advances in Covalent Kinase Inhibitors. Chem. Soc. Rev. 2020, 49 (9), 2617–2687. https://doi.org/10.1039/C9CS00720B.

(29) Strelow, J. M. A Perspective on the Kinetics of Covalent and Irreversible Inhibition. SLAS Discov. 2017, 22 (1), 3–20. https://doi.org/10.1177/1087057116671509.

(30) Almaliti, J.; Alzweiri, M.; Alhindy, M.; Al-Helo, T.; Daoud, I.; Deknash, R.; Naman, C. B.; Abu-Irmaileh, B.; Bustanji, Y.; Hamad, I. Discovery of Novel Epoxyketone Peptides as Lipase Inhibitors. Molecules 2022, 27 (7), 2261. https://doi.org/10.3390/molecules27072261.

(31) Joyce, J. A.; Baruch, A.; Chehade, K.; Meyer-Morse, N.; Giraudo, E.; Tsai, F.-Y.; Greenbaum, D. C.; Hager, J. H.; Bogyo, M.; Hanahan, D. Cathepsin Cysteine Proteases Are Effectors of Invasive Growth and Angiogenesis during Multistage Tumorigenesis. Cancer Cell 2004, 5 (5), 443–453. https://doi.org/10.1016/S1535-6108(04)00111-4.

