## Supplemental Methods, Figures and Tables for "Covalent macrocyclic proteasome inhibitors mitigate resistance in *Plasmodium falciparum*"

**Table Of Contents**

Supplemental Figure 1: HFF Cell Viability for Compounds 1-6

Supplemental Figure 2: HFF Cell Viability for Compounds 7-12

Supplemental Figure 3: HFF Cell Viability for Compounds 13-24

Supplemental Figure 4:HFF Cell Viability for Compounds 25-28, 8304-vs

Supplemental Figure 5: Pulse Assays for Compound 27,28, 8304-vs

Supplemental Figure 6: Minimum inoculum of resistance (MIR) selection experiments for 8304-vs

Supplemental Figure 7: Minimum inoculum of resistance (MIR) selection experiments for Compound 27

Supplemental Figure 8: Profiling of 20S β5 and β6 subunit mutants against 27

Supplemental Figure 9: Microsome Stability of 8304-epoxy.

General procedure for synthesis of Compounds

Supplemental Scheme 1

Supplemental Scheme 2

Supplemental Scheme 3

Supplemental Scheme 4

Supplemental Scheme 5

Mass Spectrometry Data for Compounds

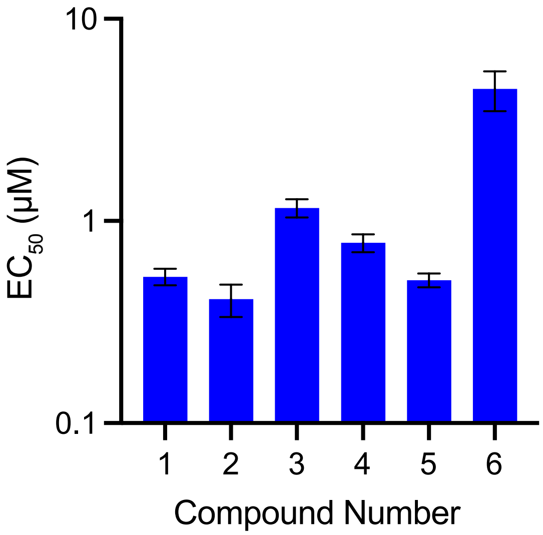


**Supplemental Figure 1**: HFF Toxicity for alkyl cyclic peptide inhibitor and linear counterparts. Compound mean ± SEM $EC_{50}$ values of each inhibitor for HFF growth. Data were generated with 72 hr assays (N, n = 2,3).

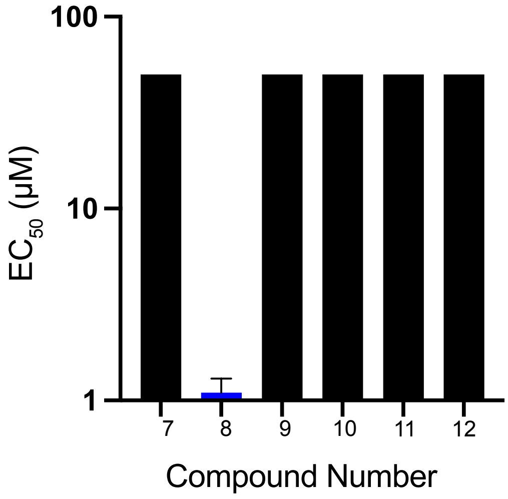


**Supplemental Figure 2**: HFF Toxicity for ether linked cyclic peptide inhibitor and linear counterparts. Compound mean ± SEM $EC_{50}$ values of each inhibitor for HFF growth. Data were generated with 72 hr assays (N, n = 2,3), Black bars denote cytotoxicity above the solubility of the compound.

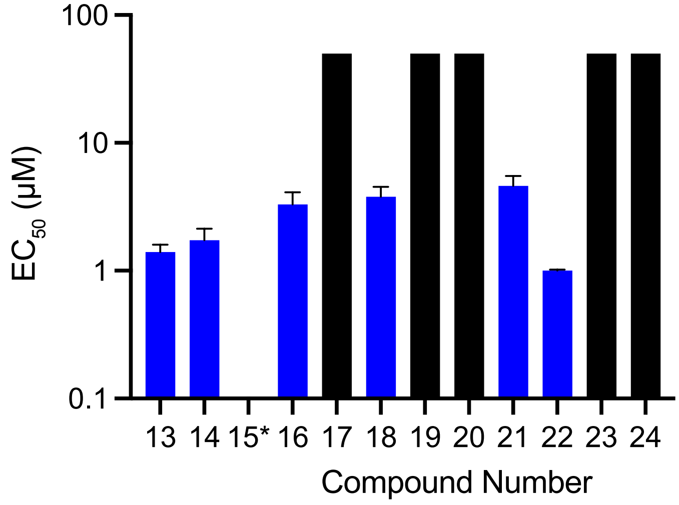


**Supplemental Figure 3**: HFF Toxicity for imidazole/pyrazol capped cyclic peptide inhibitor and linear counterparts. Compound mean ± SEM $EC_{50}$ values of each inhibitor for HFF growth. Data were generated with 72 hr assays (N, n = 2,3). Black bars denote cytotoxicity above the solubility of the compound.

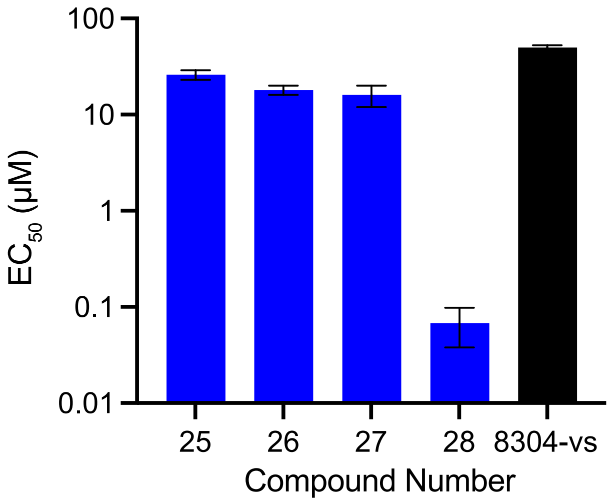


**Supplemental Figure 4**: HFF Toxicity for final set of optimized compounds. Compound mean ± SEM $EC_{50}$ values of each inhibitor for HFF growth. Data were generated with 72 hr assays (N, n = 2,3). Black bars denote cytotoxicity above the solubility of the compound.

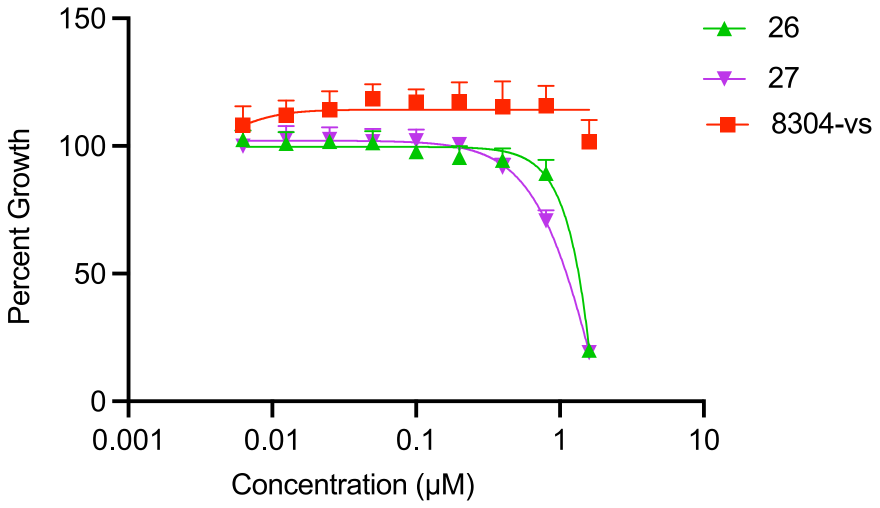


**Supplemental Figure 5**: Pulse assays for Compounds 26, 27, and 8304-vs. Killing curves with mean percent growth ± SEM for each point are shown for *P. falciparum* asexual blood stage W2 parasites, treating for one hour with each inhibitor, washed, and grown for a subsequent 72 hr. Data were generated with 72 hr assays (N, n = 6,2).

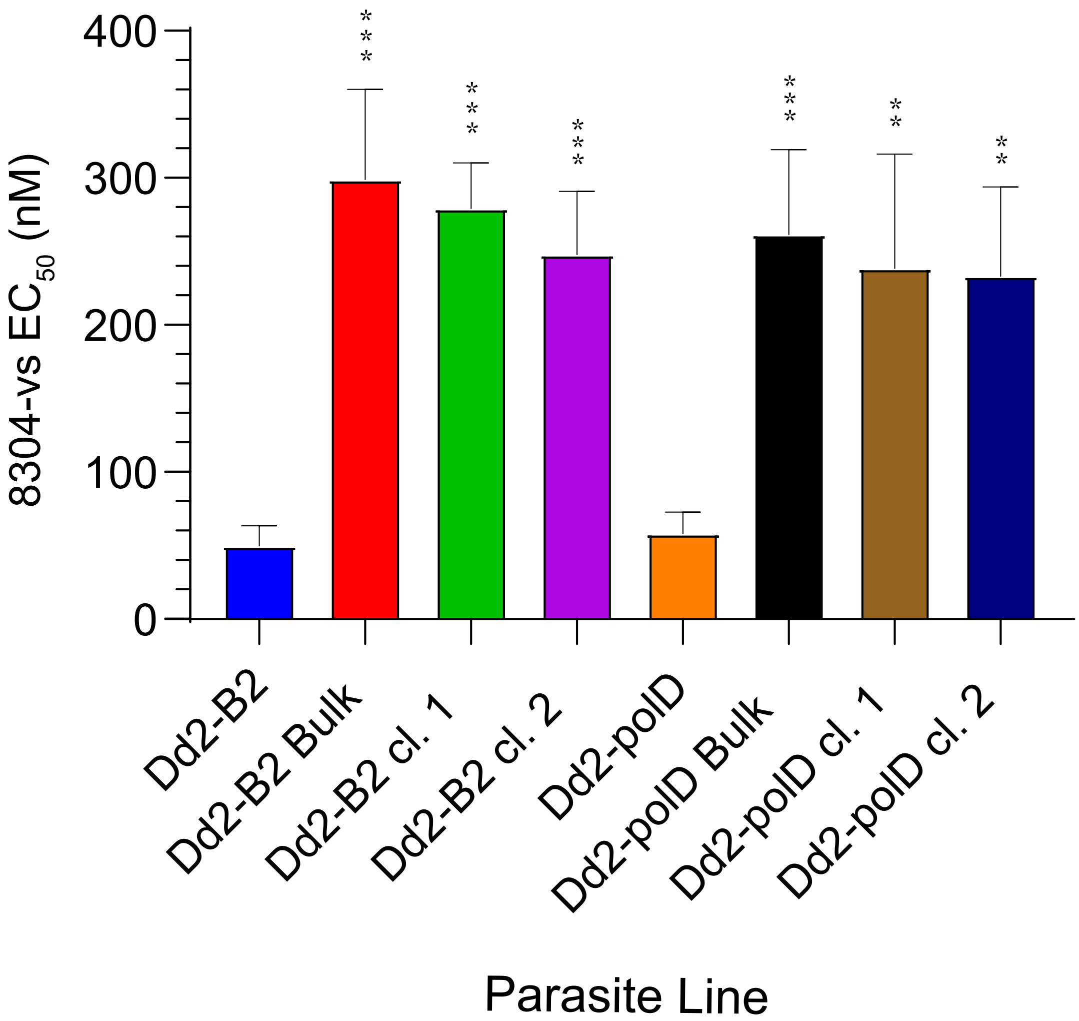


**Supplemental Figure 6.** Minimum inoculum of resistance (MIR) selection experiments for 8304-vs. Mean $IC_{50}$ values are shown for bulk cultures and clones selected for with 8304-vs. Statistical significance, compared to Dd2-B2 or Dd2-Polδ, was calculated using unpaired t tests with Welch's correction. **$p<0.01$; ***$p<0.001$. N,n = 4-5,2.

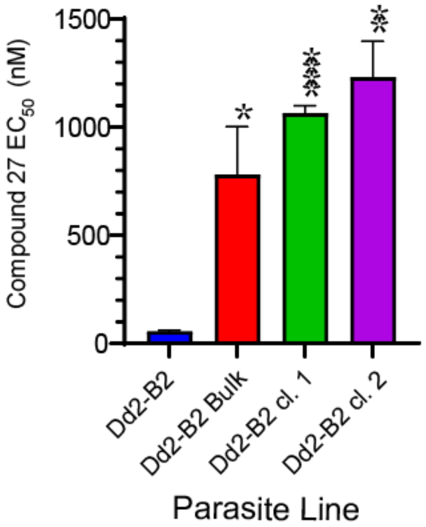


**Supplemental Figure 7.** Minimum inoculum of resistance (MIR) selection experiments for Compound 27. Mean $IC_{50}$ values are shown for bulk cultures and clones selected for with **27**. Statistical significance, compared to Dd2-B2, was calculated using unpaired t tests with Welch's correction. *p<0.05; **p<0.01; ****p<0.0001. N,n = 4-5,2.

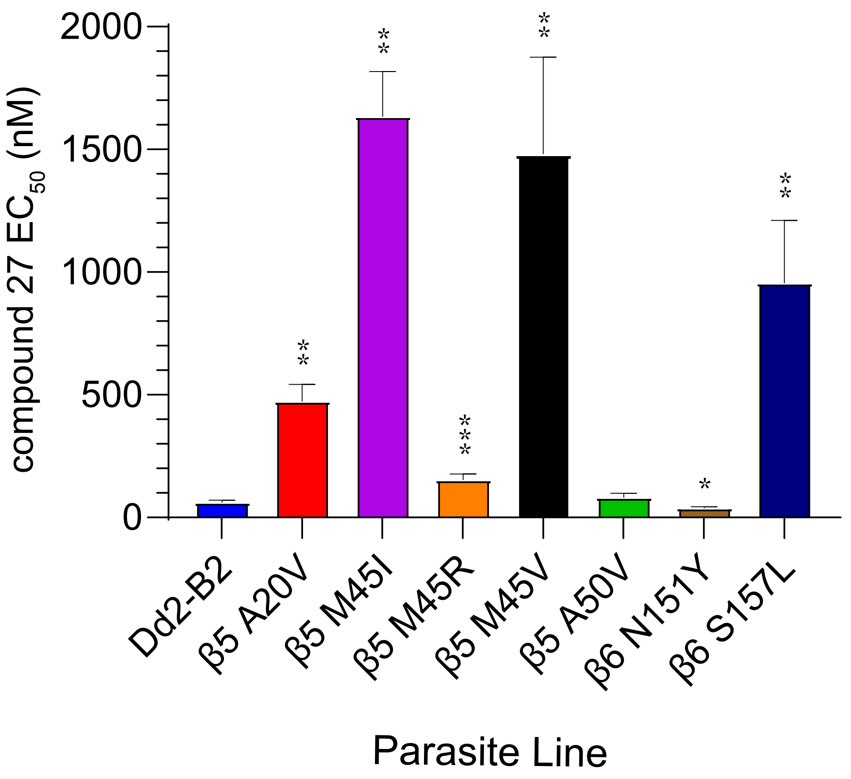


**Supplemental Figure 8:** Profiling of 20S β5 and β6 subunit mutants against 27. Dd2-B2 parasites containing previously selected proteasome mutations were assayed against compound 27. Statistical significance, compared to Dd2-B2, was calculated using unpaired t tests with Welch's correction. *p<0.05; **p<0.01; ***p<0.001. N,n = 5,2.

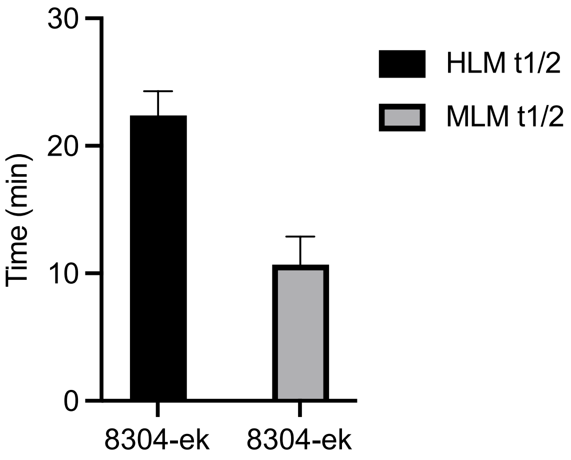


**Supplemental Figure 9:** Bar graphs depicting the half-life ± SEM of 8304-epoxy in either human (black) or mice (gray) microsomes, compound metabolism was measured by LC/MS/MS, N=2.

**General chemical materials and methods**

All solvents were purchased from Fisher Scientific (HPLC grade). All reagents were purchased from Sigma Aldrich, Chem-Impex International, and were used without further purification. All water-sensitive reactions were performed in anhydrous solvents under positive pressure of argon. Reactions were analyzed by LC-MS using an API 150EX single-quadrupole mass spectrometer (Applied Biosystems). Reverse-phase HPLC was conducted with an Agilent 1260 infinity using C18 columns (5 μm, 250 × 10 mm). NMR spectra were recorded on a Varian 400 MHz (400/100), Varian 500 MHz (500/125) or a Varian Inova 600 MHz (600/150 MHz) equipped with a pulsed field gradient accessory. Chemical shifts are given in ppm (δ) relative to tetramethylsilane as an internal standard. Coupling constants are given in Hz.

Compounds 1-6 were synthesized via Solid State Peptide Synthesis with general procedure as follows and were prepared as described previously.[1]

**General Procedures for Synthesis**

(A) Boc-Deprotections: 1 eq of Boc-protected amino acid or peptide was added to a flask and dissolved in 4M HCl in Dioxanes (0.15 M) and stirred at room temperature for 1-4 hrs. Reactions were monitored by LC/MS for completion. Solvent was then removed under reduced pressure and the product was carried forward without further purification.

(B) Capping Group Amide Couple: 1.2 eq of capping group acid was dissolved in dichloromethane (0.1 M). 5 eq of collidine was added and then 3 eq of HBTU. Lastly 1 eq of HCl Salt of dipeptide was added, and stirred at room temperature overnight. The reaction was then diluted with ethyl acetate and water (1:1 mixture). The aqueous phase was extracted twice with ethyl acetate. The combined organic was washed with saturated sodium bicarbonate solution and brine. The organic was then dried over magnesium sulfate, filtered, and concentrated under reduced pressure. The resulting residue was purified using flash chromatography.

(C) Ring Closing Metathesis: 1 eq of peptide was dissolved in dichloroethane (0.02 M) and the solution was degassed with argon. 0.05 eq of Hoveyda-Grubbs Catalyst® M720 was dissolved in 1 mL of dichloroethane and added to dissolved peptide. The reaction was stirred overnight at 40˚C. The reaction was concentrated under reduced pressure and residue was purified on a C18 reverse phase silica column (40%-90% Acetonitrile:Water + 0.1% trifluoroacetic acid).

(D) Deprotection of Benzyl and Reduction of Alkene: 1 eq of cyclized peptide was dissolved in methanol (0.2 M) and 0.1 eq of 10% palladium on carbon was added. The solution was purged with argon and then hydrogen was bubbled through and stirred for 1-16 hrs. The reaction was monitored by LC/MS for completion. Once full conversion to product, the reaction was filtered through Celite, concentrated under reduced pressure. The product was carried forward without further purification.

(E) Deprotection of Acid: 1 eq of cyclized peptide was dissolved in THF (0.2M) and 1.5 eq of LiOH was added to solution. The reaction was allowed to stir at room temperature for 1-3 hrs. The reaction was monitored by LC/MS for completion. Once full conversion to product, the reaction was acidified to pH 2 using 1 M hydrochloric acid, and then extracted three times using ethyl acetate. The combined organic was washed with brine, dried over magnesium sulfate, filtered, and concentrated under reduced pressure. The product was carried forward without further purification.

(F) Peptide Couplings: 1 eq of acid was dissolved in DMF (0.2 M) and 3 eq of HBTU, 5 eq of collidine, 1.2 eq of HCl salt amine were added and the reaction was stirred overnight. The reaction was then diluted with ethyl acetate and water (1:1 mixture). The aqueous phase was extracted twice with ethyl acetate. The combined organic was washed with saturated sodium bicarbonate solution and brine. The organic was then dried over magnesium sulfate, filtered, and concentrated under reduced pressure. The resulting residue was purified using flash chromatography.

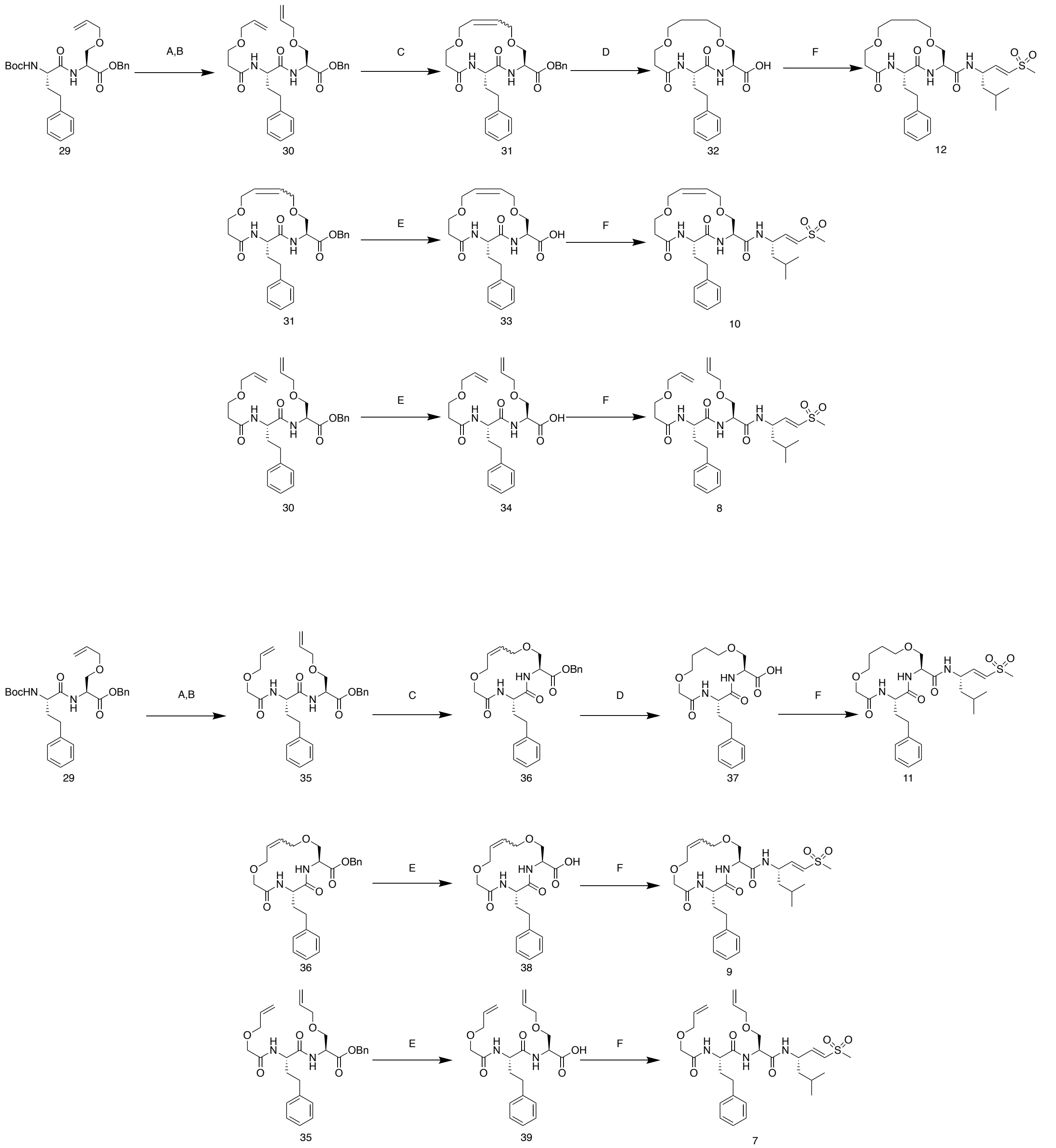


**Supplemental Scheme 1**

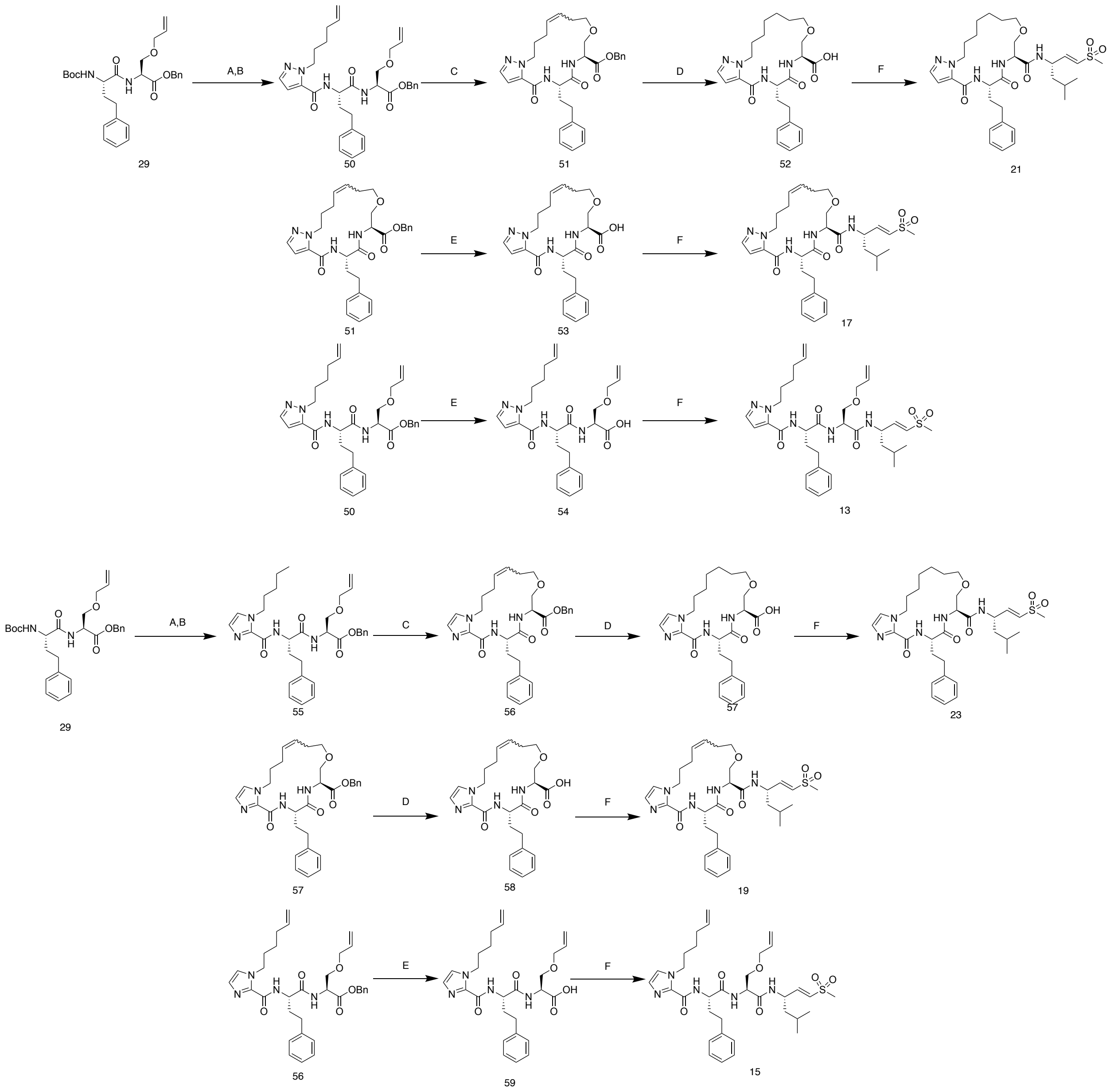


#### Supplemental Scheme 2

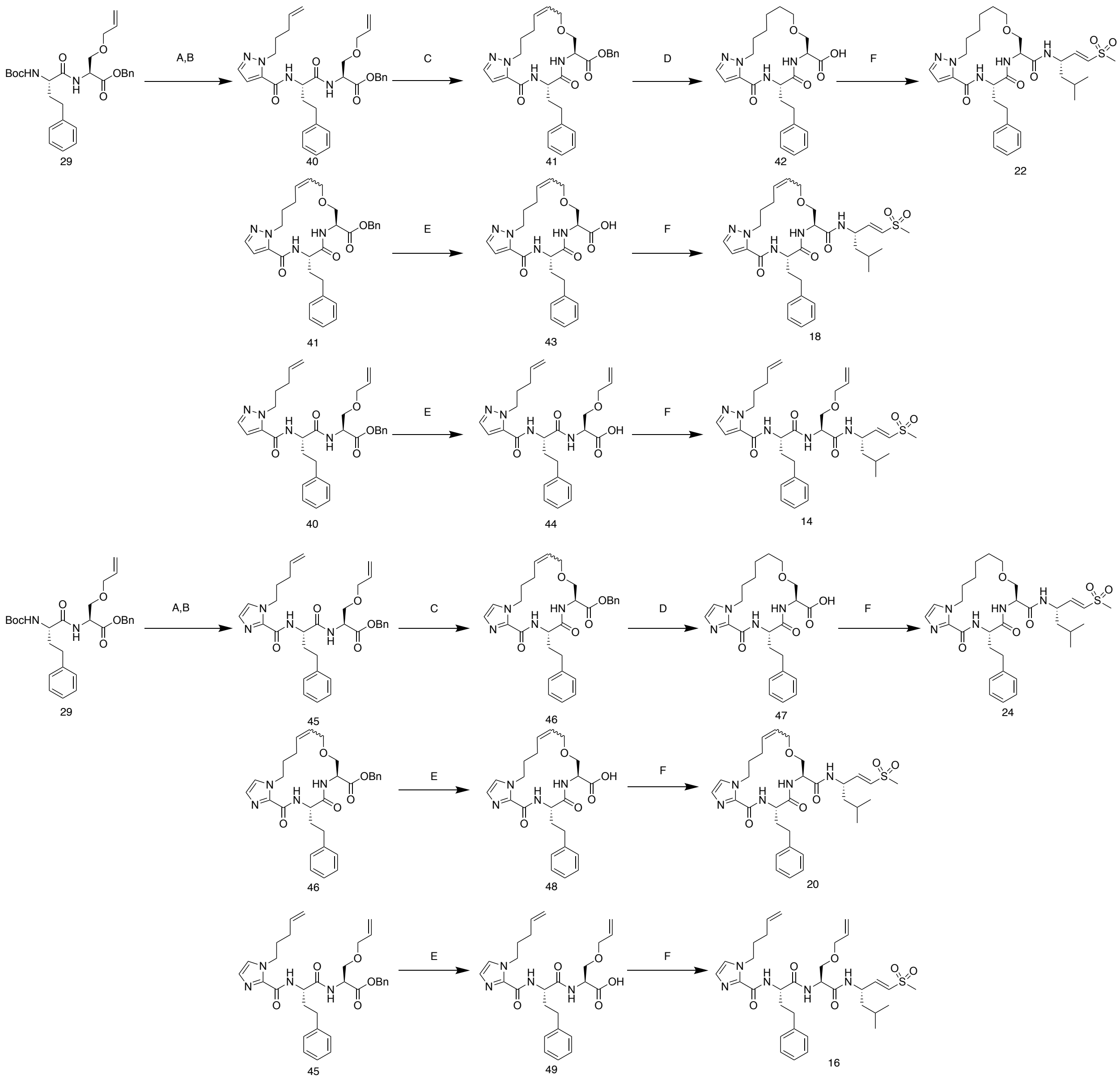


### Supplemental Scheme 3

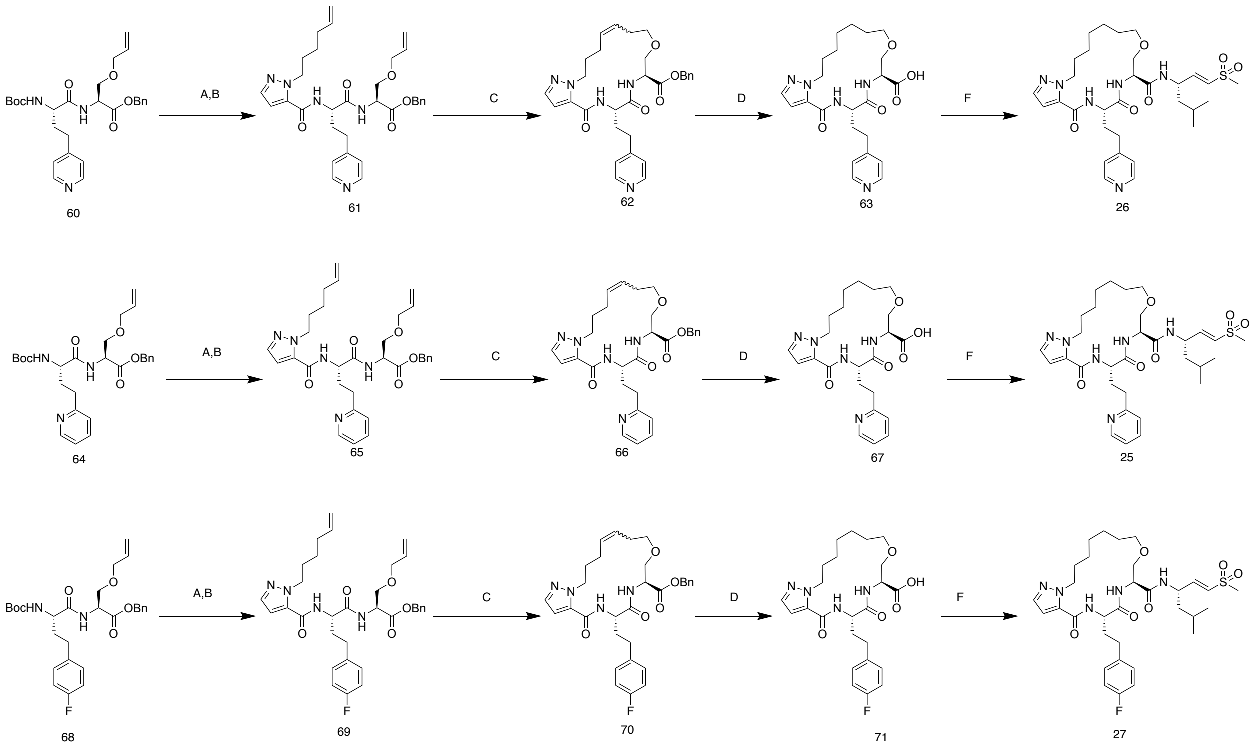


### Supplemental Scheme 4

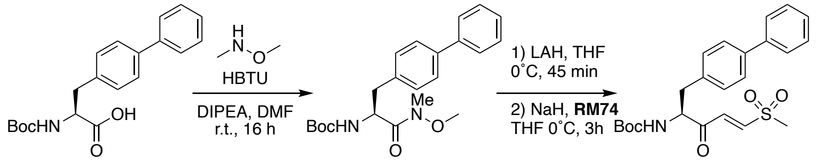


RM01-067
1.15g, 96%

72

Exact Mass: 429.16
~ 70%
73

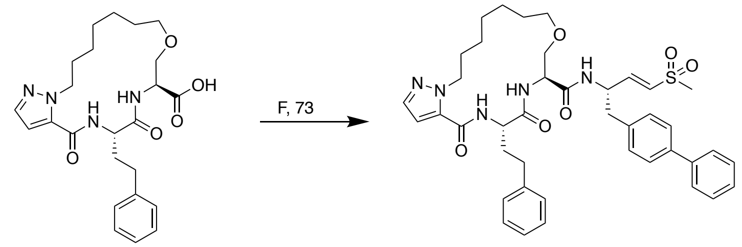


57

28

**Supplemental Scheme 5**

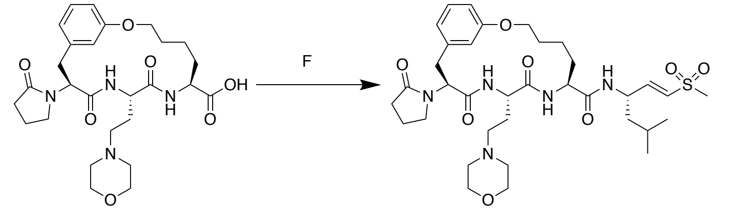


8304-vs

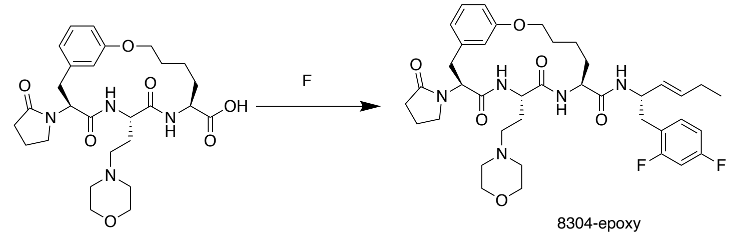


8304-epoxy

**Supplemental Scheme 6**

**Compound Mass Analysis:**

(*S*)-2-((*S*)-2-(hex-5-enamido)-4-phenylbutanamido)-*N*-((*S*,*E*)-5-methyl-1-(methylsulfonyl)hex-1-en-3-yl)hept-6-enamide (1): Exact Mass = 573.79, Found Mass[M+1] = 574.6

*N*-((*S*)-1-(((*S*)-3-(allyloxy)-1-(((*S*,*E*)-5-methyl-1-(methylsulfonyl)hex-1-en-3-yl)amino)-1-oxopropan-2-yl)amino)-1-oxo-4-phenylbutan-2-yl)hex-5-enamide (2): Exact Mass = 575.77, Found Mass [M+1]= 576.9

(*S*)-2-((*S*)-2-(hept-6-enamido)-4-phenylbutanamido)-*N*-((*S*,*E*)-5-methyl-1-(methylsulfonyl)hex-1-en-3-yl)hept-6-enamide(3): Exact Mass= 587.82, Found Mass [M+1]= 588.6

(2S,5S,Z)-N-((S,E)-5-methyl-1-(methylsulfonyl)hex-1-en-3-yl)-3,14-dioxo-2-phenethyl-1,4-diazacyclotetradec-9-ene-5-carboxamide(4): Exact Mass= 545.74, Found Mass [M+1] = 546.6

(3*S*,6*S*,*Z*)-*N*-((*S*,*E*)-5-methyl-1-(methylsulfonyl)hex-1-en-3-yl)-5,8-dioxo-6-phenethyl-1-oxa-4,7-diazacyclotetradec-12-ene-3-carboxamide(5): Exact Mass = 547.71, Found Mass [M+1] 548.4

(2*S*,5*S*,*Z*)-*N*-((*S*,*E*)-5-methyl-1-(methylsulfonyl)hex-1-en-3-yl)-3,15-dioxo-2-phenethyl-1,4-diazacyclopentadec-9-ene-5-carboxamide(6)= Exact Mass: 559.31, Found Mass [M+1]= 560.6

(*S*)-*N*-((*S*)-3-(allyloxy)-1-(((*S*,*E*)-5-methyl-1-(methylsulfonyl)hex-1-en-3-yl)amino)-1-oxopropan-2-yl)-2-(2-(allyloxy)acetamido)-4-phenylbutanamide(7): Exact Mass= 577.28, Found Mass [M+1]= 578.3

(*S*)-*N*-((*S*)-3-(allyloxy)-1-(((*S*,*E*)-5-methyl-1-(methylsulfonyl)hex-1-en-3-yl)amino)-1-oxopropan-2-yl)-2-(3-(allyloxy)propanamido)-4-phenylbutanamide (8): Exact Mass = 591.30, Found Mass [M+1]= 592.4

(3*S*,6*S*)-*N*-((*S*,*E*)-5-methyl-1-(methylsulfonyl)hex-1-en-3-yl)-5,8-dioxo-6-phenethyl-1,10-dioxa-4,7-diazacyclotetradec-12-ene-3-carboxamide (9): Exact Mass= 549.25, Found Mass [M+1]= 550.4

(3*S*,6*S*,*Z*)-*N*-((*S*,*E*)-5-methyl-1-(methylsulfonyl)hex-1-en-3-yl)-5,8-dioxo-6-phenethyl-1,11-dioxa-4,7-diazacyclopentadec-13-ene-3-carboxamide (10): Exact Mass= 563.27, Found Mass [M+1]= 564.4

(3*S*,6*S*)-*N*-((*S*,*E*)-5-methyl-1-(methylsulfonyl)hex-1-en-3-yl)-5,8-dioxo-6-phenethyl-1,10-dioxa-4,7-diazacyclotetradecane-3-carboxamide (11): Exact Mass = 551.27, Found Mass [M+1]= 552.5

(3*S*,6*S*)-*N*-((*S*,*E*)-5-methyl-1-(methylsulfonyl)hex-1-en-3-yl)-5,8-dioxo-6-phenethyl-1,11-dioxa-4,7-diazacyclopentadecane-3-carboxamide (12): Exact Mass = 565.28, Found Mass[M+1]= 566.5

*N*-((*S*)-1-(((*S*)-3-(allyloxy)-1-(((*S*,*E*)-5-methyl-1-(methylsulfonyl)hex-1-en-3-yl)amino)-1-oxopropan-2-yl)amino)-1-oxo-4-phenylbutan-2-yl)-1-(hex-5-en-1-yl)-1*H*-pyrazole-5-carboxamide (13): Exact Mass = 641.32, Found Mass[M+1]= 542.2

*N*-((*S*)-1-(((*S*)-3-(allyloxy)-1-(((*S*,*E*)-5-methyl-1-(methylsulfonyl)hex-1-en-3-yl)amino)-1-oxopropan-2-yl)amino)-1-oxo-4-phenylbutan-2-yl)-1-(pent-4-en-1-yl)-1*H*-pyrazole-5-carboxamide(14): Exact Mass= 655.34, Mass Found [M+1]=656.2

*N*-((*S*)-1-(((*S*)-3-(allyloxy)-1-(((*S*,*E*)-5-methyl-1-(methylsulfonyl)hex-1-en-3-yl)amino)-1-oxopropan-2-yl)amino)-1-oxo-4-phenylbutan-2-yl)-1-(pent-4-en-1-yl)-1*H*-imidazole-2-carboxamide(16): Exact Mass: 655.34, Mass Found[M+1]= 656.7

(6*S*,9*S*)-*N*-((*S*,*E*)-5-methyl-1-(methylsulfonyl)hex-1-en-3-yl)-4,7-dioxo-6-phenethyl-5,6,7,8,9,10,13,16,17,18-decahydro-4*H*,12*H*-pyrazolo[5,1-*i*][1]oxa[4,7,10]triazacycloheptadecine-9-carboxamide(17): Exact Mass= 613.29, Mass Found [M+1]=614.4

(6*S*,9*S*)-*N*-((*S*,*E*)-5-methyl-1-(methylsulfonyl)hex-1-en-3-yl)-4,7-dioxo-6-phenethyl-5,6,7,8,9,10,12,15,16,17-decahydro-4*H*-pyrazolo[5,1-*i*][1]oxa[4,7,10]triazacyclohexadecine-9-carboxamide(18): Exact Mass= 627.31, Found Mass [M+1]= 628.4

(14*S*,17*S*)-*N*-((*S*,*E*)-5-methyl-1-(methylsulfonyl)hex-1-en-3-yl)-16,19-dioxo-17-phenethyl-6,7,10,11,14,15,16,17,18,19-decahydro-5*H*,13*H*-imidazo[2,1-*i*][1]oxa[4,7,10]triazacycloheptadecine-14-carboxamide (19): Exact Mass= 613.29, Mass Found [M+1]=614.4

(13*S*,16*S*)-*N*-((*S*,*E*)-5-methyl-1-(methylsulfonyl)hex-1-en-3-yl)-15,18-dioxo-16-phenethyl-5,6,7,10,13,14,15,16,17,18-decahydro-12*H*-imidazo[2,1-*i*][1]oxa[4,7,10]triazacyclohexadecine-13-carboxamide (20): Exact Mass= 627.31, Found Mass [M+1]= 628.4

(6*S*,9*S*)-*N*-((*S*,*E*)-5-methyl-1-(methylsulfonyl)hex-1-en-3-yl)-4,7-dioxo-6-phenethyl-5,6,7,8,9,10,13,14,15,16,17,18-dodecahydro-4*H*,12*H*-pyrazolo[5,1-*i*][1]oxa[4,7,10]triazacycloheptadecine-9-carboxamide (21): Exact Mass: 615.31, Found Mass[M+1]= 616.1

(6*S*,9*S*)-*N*-((*S*,*E*)-5-methyl-1-(methylsulfonyl)hex-1-en-3-yl)-4,7-dioxo-6-phenethyl-5,6,7,8,9,10,12,13,14,15,16,17-dodecahydro-4*H*-pyrazolo[5,1-*i*][1]oxa[4,7,10]triazacyclohexadecine-9-carboxamide (22): Exact Mass: 629.32, Found Mass [M+1]= 630.4

(14*S*,17*S*)-*N*-((*S*,*E*)-5-methyl-1-(methylsulfonyl)hex-1-en-3-yl)-16,19-dioxo-17-phenethyl-6,7,8,9,10,11,14,15,16,17,18,19-dodecahydro-5*H*,13*H*-imidazo[2,1-*i*][1]oxa[4,7,10]triazacycloheptadecine-14-carboxamide (23): Exact Mass: 615.31, Found Mass[M+1]= 616.3

(13*S*,16*S*)-*N*-((*S*,*E*)-5-methyl-1-(methylsulfonyl)hex-1-en-3-yl)-15,18-dioxo-16-phenethyl-5,6,7,8,9,10,13,14,15,16,17,18-dodecahydro-12*H*-imidazo[2,1-*i*][1]oxa[4,7,10]triazacyclohexadecine-13-carboxamide(24): act Mass: 629.32, Found Mass [M+1]= 630.6

(6*S*,9*S*)-*N*-((*S*,*E*)-5-methyl-1-(methylsulfonyl)hex-1-en-3-yl)-4,7-dioxo-6-(2-(pyridin-2-yl)ethyl)-5,6,7,8,9,10,13,14,15,16,17,18-dodecahydro-4*H*,12*H*-pyrazolo[5,1-*i*][1]oxa[4,7,10]triazacycloheptadecine-9-carboxamide(25): Exact Mass: 630.32, Found Mass[M+1]= 631.3

(6*S*,9*S*)-*N*-((*S*,*E*)-5-methyl-1-(methylsulfonyl)hex-1-en-3-yl)-4,7-dioxo-6-(2-(pyridin-4-yl)ethyl)-5,6,7,8,9,10,13,14,15,16,17,18-dodecahydro-4*H*,12*H*-pyrazolo[5,1-*i*][1]oxa[4,7,10]triazacycloheptadecine-9-carboxamide (26): Exact Mass: 630.32, Found Mass[M+1]=631.7

(6*S*,9*S*)-6-(4-fluorophenethyl)-*N*-((*S*,*E*)-5-methyl-1-(methylsulfonyl)hex-1-en-3-yl)-4,7-dioxo-5,6,7,8,9,10,13,14,15,16,17,18-dodecahydro-4*H*,12*H*-pyrazolo[5,1-*i*][1]oxa[4,7,10]triazacycloheptadecine-9-carboxamide (27): Exact Mass: 647.32, Found Mass[M+1]= 648.3

(6*S*,9*S*)-*N*-((*S*,*E*)-1-([1,1'-biphenyl]-4-yl)-4-(methylsulfonyl)but-3-en-2-yl)-4,7-dioxo-6-phenethyl-5,6,7,8,9,10,13,14,15,16,17,18-dodecahydro-4*H*,12*H*-pyrazolo[5,1-*i*][1]oxa[4,7,10]triazacycloheptadecine-9-carboxamide (28): Exact Mass: 739.34, Found Mass[M+1]=740.5

(7*S*,10*S*,13*S*)-*N*-((*S*,*E*)-5-methyl-1-(methylsulfonyl)hex-1-en-3-yl)-10-(2-morpholinoethyl)-9,12-dioxo-13-(2-oxopyrrolidin-1-yl)-2-oxa-8,11-diaza-1(1,3)-benzenacyclotetradecaphane-7-carboxamide (8304-vs): Exact Mass: 703.36, Found Mass = 704.4

(7*S*,10*S*,13*S*)-*N*-((*S*,*E*)-1-(2,4-difluorophenyl)hex-3-en-2-yl)-10-(2-morpholinoethyl)-9,12-dioxo-13-(2-oxopyrrolidin-1-yl)-2-oxa-8,11-diaza-1(1,3)-benzenacyclotetradecaphane-7-carboxamide (8304-epoxy): Exact Mass: 753.35, Found Mass[M+H]= 754.4

benzyl *O*-allyl-*N*-((*S*)-2-((*tert*-butoxycarbonyl)amino)-4-phenylbutanoyl)-*L*-serinate (29): Calculated Mass: 496.26. Found Mass[M-1]: 495.0

benzyl *O*-allyl-*N*-((*S*)-2-(3-(allyloxy)propanamido)-4-phenylbutanoyl)-*L*-serinate (30): Calculated Mass: 508.26 Found Mass[M+1]: 509.3

benzyl (3*S*,6*S*)-5,8-dioxo-6-phenethyl-1,11-dioxa-4,7-diazacyclopentadec-13-ene-3-carboxylate (31):

Exact Mass: 480.23, Found Mass[M+1]: 481.1

(3*S*,6*S*)-5,8-dioxo-6-phenethyl-1,11-dioxa-4,7-diazacyclopentadecane-3-carboxylic acid (32): Exact Mass: 392.19, Found Mass[M+1]: 393.4

(3*S*,6*S*,*Z*)-5,8-dioxo-6-phenethyl-1,11-dioxa-4,7-diazacyclopentadec-13-ene-3-carboxylic acid (33): Exact Mass: 390.18, Found Mass[M+1]= 391.4

*O*-allyl-*N*-((*S*)-2-(3-(allyloxy)propanamido)-4-phenylbutanoyl)-*L*-serine (34): Exact Mass: 418.21, Found Mass[M+1]= 419.6

benzyl *O*-allyl-*N*-((*S*)-2-(2-(allyloxy)acetamido)-4-phenylbutanoyl)-*L*-serinate (35): Exact Mass: 494.24, Found Mass [M+1]= 495.1

benzyl (3*S*,6*S*)-5,8-dioxo-6-phenethyl-1,10-dioxa-4,7-diazacyclotetradec-12-ene-3-carboxylate (36): Exact Mass: 466.21, Found Mass[M+1]= 467.1

(3*S*,6*S*)-5,8-dioxo-6-phenethyl-1,10-dioxa-4,7-diazacyclotetradecane-3-carboxylic acid (37): Exact Mass: 378.18, Found Mass[M+1]= 379.3

(3*S*,6*S*)-5,8-dioxo-6-phenethyl-1,10-dioxa-4,7-diazacyclotetradec-12-ene-3-carboxylic acid (38): Exact Mass: 376.16, Found Mass[M+1]= 377.2

*O*-allyl-*N*-((*S*)-2-(2-(allyloxy)acetamido)-4-phenylbutanoyl)-*L*-serine (39): Exact Mass: 404.19, Found Mass[M+1]=405.2

benzyl *O*-allyl-*N*-((*S*)-2-(1-(pent-4-en-1-yl)-1*H*-pyrazole-5-carboxamido)-4-phenylbutanoyl)-*L*-serinate (40): Exact Mass: 558.28, Found Mass[M+1]= 559.3

benzyl (6*S*,9*S*)-4,7-dioxo-6-phenethyl-5,6,7,8,9,10,12,15,16,17-decahydro-4*H*-pyrazolo[5,1-*i*][1]oxa[4,7,10]triazacyclohexadecine-9-carboxylate (41): Exact Mass: 530.25, Found Mass[M+1]: 531.4

(6*S*,9*S*)-4,7-dioxo-6-phenethyl-5,6,7,8,9,10,12,13,14,15,16,17-dodecahydro-4*H*-pyrazolo[5,1-*i*][1]oxa[4,7,10]triazacyclohexadecine-9-carboxylic acid (42): Exact Mass: 442.22, Found Mass[M+1] =443.4

(6*S*,9*S*)-4,7-dioxo-6-phenethyl-5,6,7,8,9,10,12,15,16,17-decahydro-4*H*-pyrazolo[5,1-*i*][1]oxa[4,7,10]triazacyclohexadecine-9-carboxylic acid (43): Exact Mass: 440.21, Found Mass[M+1]= 441.3

*O*-allyl-*N*-((*S*)-2-(1-(pent-4-en-1-yl)-1*H*-pyrazole-5-carboxamido)-4-phenylbutanoyl)-*L*-serine (44): Exact Mass: 468.24, Found Mass [M+1] = 469.1

benzyl *O*-allyl-*N*-((*S*)-2-(1-(pent-4-en-1-yl)-1*H*-imidazole-2-carboxamido)-4-phenylbutanoyl)-*L*-serinate (45): Exact Mass: 558.28, Found Mass [M+1] = 559.5

benzyl (13*S*,16*S*)-15,18-dioxo-16-phenethyl-5,6,7,10,13,14,15,16,17,18-decahydro-12*H*-imidazo[2,1-*i*][1]oxa[4,7,10]triazacyclohexadecine-13-carboxylate (46): Exact Mass: 530.25, Found Mass [M+1]= 531.4

(13*S*,16*S*)-15,18-dioxo-16-phenethyl-5,6,7,8,9,10,13,14,15,16,17,18-dodecahydro-12*H*-imidazo[2,1-*i*][1]oxa[4,7,10]triazacyclohexadecine-13-carboxylic acid (47): Exact Mass: 442.22, Found Mass [M+1]= 443.5

(13*S*,16*S*)-15,18-dioxo-16-phenethyl-5,6,7,10,13,14,15,16,17,18-decahydro-12*H*-imidazo[2,1-*i*][1]oxa[4,7,10]triazacyclohexadecine-13-carboxylic acid (48): Exact Mass: 440.21, Found Mass [M+1]=441.3

*O*-allyl-*N*-((*S*)-2-(1-(pent-4-en-1-yl)-1*H*-imidazole-2-carboxamido)-4-phenylbutanoyl)-*L*-serine (49): Exact Mass: 468.24, Found Mass [M+1] = 496.5

benzyl *O*-allyl-*N*-((*S*)-2-(1-(hex-5-en-1-yl)-1*H*-pyrazole-5-carboxamido)-4-phenylbutanoyl)-*L*-serinate (50): Exact Mass: 572.30, Found Mass[M+1] = 572.9

benzyl (6*S*,9*S*)-4,7-dioxo-6-phenethyl-5,6,7,8,9,10,13,16,17,18-decahydro-4*H*,12*H*-pyrazolo[5,1-*i*][1]oxa[4,7,10]triazacycloheptadecine-9-carboxylate (51): Exact Mass: 544.27, Found Mass [M+1]: 545.0

(6*S*,9*S*)-4,7-dioxo-6-phenethyl-5,6,7,8,9,10,13,14,15,16,17,18-dodecahydro-4*H*,12*H*-pyrazolo[5,1-*i*][1]oxa[4,7,10]triazacycloheptadecine-9-carboxylic acid (52): Exact Mass: 456.24, Found Maxx [M+1] = 457.2

(6*S*,9*S*)-4,7-dioxo-6-phenethyl-5,6,7,8,9,10,13,16,17,18-decahydro-4*H*,12*H*-pyrazolo[5,1-*i*][1]oxa[4,7,10]triazacycloheptadecine-9-carboxylic acid (53): Exact Mass: 454.22, Found Mass [M+1]: 455.1

*O*-allyl-*N*-((*S*)-2-(1-(hex-5-en-1-yl)-1*H*-pyrazole-5-carboxamido)-4-phenylbutanoyl)-*L*-serine (54): Exact Mass: 482.25, Found Mass [M+1] = 483.2

benzyl *O*-allyl-*N*-((*S*)-2-(1-pentyl-1*H*-imidazole-2-carboxamido)-4-phenylbutanoyl)-*L*-serinate (55): Exact Mass: 572.3, Found Mass [M +1] = 572.9

benzyl (14*S*,17*S*)-16,19-dioxo-17-phenethyl-6,7,10,11,14,15,16,17,18,19-decahydro-5*H*,13*H*-imidazo[2,1-*i*][1]oxa[4,7,10]triazacycloheptadecine-14-carboxylate(56): Exact Mass: 544.27, Found Mass [M+1] = 545.5

(14*S*,17*S*)-16,19-dioxo-17-phenethyl-6,7,8,9,10,11,14,15,16,17,18,19-dodecahydro-5*H*,13*H*-imidazo[2,1-*i*][1]oxa[4,7,10]triazacycloheptadecine-14-carboxylic acid (57): Exact Mass: 456.24, Found Mass [M+1] = 457.2

(14*S*,17*S*)-16,19-dioxo-17-phenethyl-6,7,10,11,14,15,16,17,18,19-decahydro-5*H*,13*H*-imidazo[2,1-*i*][1]oxa[4,7,10]triazacycloheptadecine-14-carboxylic acid (58): Exact Mass: 454.22, Found Mass [M+1]= 455.4

*O*-allyl-*N*-((*S*)-2-(1-(hex-5-en-1-yl)-1*H*-imidazole-2-carboxamido)-4-phenylbutanoyl)-*L*-serine (59): Exact Mass: 482.25, Found Mass [M+1]= 483.3

benzyl *O*-allyl-*N*-((*S*)-2-((*tert*-butoxycarbonyl)amino)-4-(pyridin-4-yl)butanoyl)-*L*-serinate (60): Exact Mass: 497.25, Found Mass [M+1]= 498.4

benzyl *O*-allyl-*N*-((*S*)-2-(1-(hex-5-en-1-yl)-1*H*-pyrazole-5-carboxamido)-4-(pyridin-4-yl)butanoyl)-*L*-serinate (61): Exact Mass: 573.30, Fond Mass [M+]= 573.3

benzyl (6*S*,9*S*)-4,7-dioxo-6-(2-(pyridin-4-yl)ethyl)-5,6,7,8,9,10,13,16,17,18-decahydro-4*H*,12*H*-pyrazolo[5,1-*i*][1]oxa[4,7,10]triazacycloheptadecine-9-carboxylate (62): Exact Mass: 545.26, Found Mass [M+1]= 546.3

(6*S*,9*S*)-4,7-dioxo-6-(2-(pyridin-4-yl)ethyl)-5,6,7,8,9,10,13,14,15,16,17,18-dodecahydro-4*H*,12*H*-pyrazolo[5,1-*i*][1]oxa[4,7,10]triazacycloheptadecine-9-carboxylic acid (63): Exact Mass: 457.23, Found Mass [M-1]= 456.2

benzyl *O*-allyl-*N*-((*S*)-2-((*tert*-butoxycarbonyl)amino)-4-(pyridin-2-yl)butanoyl)-*L*-serinate (64): Exact Mass: 497.25, Found Mass [M+1]= 498.4

benzyl *O*-allyl-*N*-((*S*)-2-(1-(hex-5-en-1-yl)-1*H*-pyrazole-5-carboxamido)-4-(pyridin-2-yl)butanoyl)-*L*-serinate (65): Exact Mass: 573.30, Found Mass [M+1]= 574.3

benzyl (6*S*,9*S*)-4,7-dioxo-6-(2-(pyridin-2-yl)ethyl)-5,6,7,8,9,10,13,16,17,18-decahydro-4*H*,12*H*-pyrazolo[5,1-*i*][1]oxa[4,7,10]triazacycloheptadecine-9-carboxylate (66): Exact Mass: 545.26, Found Mass [M+1]= 546.4

(6*S*,9*S*)-4,7-dioxo-6-(2-(pyridin-2-yl)ethyl)-5,6,7,8,9,10,13,14,15,16,17,18-dodecahydro-4*H*,12*H*-pyrazolo[5,1-*i*][1]oxa[4,7,10]triazacycloheptadecine-9-carboxylic acid (67): Exact Mass: 457.23, Found Mass [M-1]= 456.2

benzyl *O*-allyl-*N*-((*S*)-2-((*tert*-butoxycarbonyl)amino)-4-(4-fluorophenyl)butanoyl)-*L*-serinate (68): Exact Mass: 514.25, Found Mass[M+1]=515.3

benzyl *O*-allyl-*N*-((*S*)-4-(4-fluorophenyl)-2-(1-(hex-5-en-1-yl)-1*H*-pyrazole-5-carboxamido)butanoyl)-*L*-serinate (69): Exact Mass: 590.29, Found Mass [M+1]= 591.4

benzyl (6*S*,9*S*)-6-(4-fluorophenethyl)-4,7-dioxo-5,6,7,8,9,10,13,16,17,18-decahydro-4*H*,12*H*-pyrazolo[5,1-*i*][1]oxa[4,7,10]triazacycloheptadecine-9-carboxylate (70): Exact Mass: 562.26, Found Mass [M+1]= 563.3

(6*S*,9*S*)-6-(4-fluorophenethyl)-4,7-dioxo-5,6,7,8,9,10,13,14,15,16,17,18-dodecahydro-4*H*,12*H*-pyrazolo[5,1-*i*][1]oxa[4,7,10]triazacycloheptadecine-9-carboxylic acid (71): Exact Mass: 474.23, Found Mass [M-1]=473.3

*tert*-butyl (*S*)-(3-([1,1'-biphenyl]-4-yl)-1-(methoxy(methyl)amino)-1-oxopropan-2-yl)carbamate (72): Exact Mass: 384.20, Found Mass [M-1]=383.2

*tert*-butyl (*S*,*E*)-(1-([1,1'-biphenyl]-4-yl)-5-(methylsulfonyl)-3-oxopent-4-en-2-yl)carbamate (73): $^{1}$H NMR (500 MHz, MeOD) δ 7.69 – 7.62 (m, 5H), 7.48 – 7.36 (m, 6H), 6.87 – 6.75 (m, 2H), 4.35 (q, *J* = 7.4 Hz, 1H), 3.23 (td, *J* = 15.6, 14.7, 5.8 Hz, 1H), 3.16 – 3.03 (m, 1H), 2.93 (s, 3H).
